# Estimation of the mass density of biological matter from refractive index measurements

**DOI:** 10.1101/2023.12.05.569868

**Authors:** Conrad Möckel, Timon Beck, Sara Kaliman, Shada Abuhattum, Kyoohyun Kim, Julia Kolb, Daniel Wehner, Vasily Zaburdaev, Jochen Guck

## Abstract

The quantification of physical properties of biological matter gives rise to novel ways of understanding functional mechanisms by utilizing models that explicitly depend on physical observables. One of the basic biophysical properties is the mass density (MD), which determines the degree of crowdedness. It impacts the dynamics in sub-cellular compartments and further plays a major role in defining the opto-acoustical properties of cells and tissues. As such, the MD can be connected to the refractive index (RI) via the well known Lorentz-Lorenz relation, which takes into account the polarizability of matter. However, computing the MD based on RI measurements poses a challenge as it requires detailed knowledge of the biochemical composition of the sample. Here we propose a methodology on how to account for *a priori* and *a posteriori* assumptions about the biochemical composition of the sample as well as respective RI measurements. To that aim, we employ the Biot mixing rule of RIs alongside the assumption of volume additivity to find an approximate relation of MD and RI. We use Monte-Carlo simulations as well as Gaussian propagation of uncertainty to obtain approximate analytical solutions for the respective uncertainties of MD and RI. We validate this approach by applying it to a set of well characterized complex mixtures given by *bovine* milk and intralipid emulsion. Further, we employ it to estimate the mass density of trunk tissue of living zebrafish (*Danio rerio*) larvae. Our results enable quantifying changes of mass density estimates based on variations in the *a priori* assumptions. This illustrates the importance of implementing this methodology not only for MD estimations but for many other related biophysical problems, such as mechanical measurements using Brillouin microscopy and transient optical coherence elastography.

## I. INTRODUCTION

Quantifying the physical properties of biological matter has become increasingly important over recent decades. By now, the notion that biological function of cells and tissues is affected by their physical phenotype and vice versa has been validated in many experimental studies (see e.g., [1–3]). A fundamental property of matter, including living matter, is the mass density (MD) [4], which is not only associated with buoyancy, crowdedness [5], biomolecular condensation [6] and inherent dynamical processes of the sample of interest [7, 8], but also plays a major role in elastography, particularly Brillouin microscopy [9–12] and transient optical coherence elastography (see e.g., [13, 14]). However, measuring the *in vivo* MD distribution in a direct manner poses a challenge which has not been resolved so far. One way of inferring the *in vivo* MD of a sample is to measure the refractive index (RI) *via* microscopy techniques such as optical diffraction tomography (ODT) [15]. The Lorentz-Lorenz relation then connects the RI with the mass density if the molar refractivity and partial specific volume (PSV) of the dry mass composition as well as the solvent content are known. This knowledge, however, is not trivially obtainable. A customary assumption regarding biological matter is that the dry mass composition is given by proteins only [15–17] and that the solvent content is then indirectly constrained by the measured RI. While this approximation holds for binary polymer-solutions, it cannot be directly extended to samples with a ‘complex dry mass composition’. In the context of cells and tissues, the dry mass composition may be thought of as a mixture of (phase separated) proteins, lipids, sugars, etc. [18]. By employing e.g., mass spectrometry (MS) and/or (stimulated) Raman spectroscopy (SRS), individual components and their respective concentrations in the sample can be identified [19–21]. Additionally, correlative fluorescence information could be employed to segment RI maps acquired by ODT [12, 22]. However, these experimental modalities might not be available or applicable for certain samples, which creates a degree of ignorance about the dry mass composition which should be accounted for in the inference process of obtaining an MD estimate. Another, closely related aspect is the robust estimation of the uncertainty of the MD. Considering the previously mentioned degree of ignorance, these uncertainties are clearly not only of statistical but also of systematic nature. And further, even if universal knowledge about the true distributions of the molar refractivity and PSV was available, in order to estimate the uncertainty of the MD adequately, the uncertainties of the individual parameters should be propagated.

Here we present a robust methodology for estimating the uncertainties of the MD and the correlative RI by employing Monte-Carlo (MC) simulations. Furthermore, we provide analytical approximations for both the MD and RI distributions in dependence of the dry mass composition and the solvent content, employing Gaussian propagation of uncertainty (GPU).

To this end, we first motivate a simple mixture model to estimate the MD and the RI from two material constants, namely the RI increment *α* and the PSV *θ*. We then extend the model towards unimodal distributions of *α* and *θ*, for which previously only precise values were assumed.

The distributions of the RI increment and PSV are remarkably narrow, when only considering proteins in the mixture [23], resulting in sharp distributions of RI and MD. However, taking a second type of molecule, such as lipids or sugars, into account adds an additional complexity to the MD estimations since their values of RI increment and PSV differ drastically from those of proteins. Therefore, we derive an effective description of the system based on weighted mixture distributions. This allows for a correlative prediction of the RI and the MD, accounting e.g., for the lipid and water content of the sample as well as for fluctuations in both quantities. We then apply this approach to 1) *bovine* milk, a well characterized mixture of water, proteins and lipids and 2) 20% intralipid emulsion, which mainly consists of water and soybean oil. Comparing the measured values of the MD and RI acquired by pycnometry and Abbe refractometry, respectively, with our theoretical estimates, we find both to be in good agreement with each other.

After demonstrating the applicability of our method on *bovine* milk and intralipid emulsion, we explore the MD distribution of larval zebrafish trunk, compromising major tissue including muscle, spinal cord etc., employing the recent RI and MS measurements of [19]. This purely optical and computational approach shows how the MD can be estimated also in complex *in vivo* specimens, enabling a more profound interpretation of mechanical measurements.

## II. A BINARY MIXTURE MODEL FOR MASS DENSITY ESTIMATIONS

Considering a mixture of some molecule (i.e. solute content) in a solvent, the total mass *m* and volume *v* of the solution follow the form

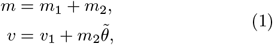

where index 1 denotes the solvent, index 2 denotes the solute and 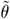 denotes the so called apparent specific volume of the solute (ASV), which describes the volume per gram of the solute *in solution*. As such, the ASV may be dependent on the mass of the solute *m*_2_, since it accounts for solute-solute interactions under constant temperature *T*, pressure *p* and solvent mass *m*_1_. The change of the total volume of the solution *v* with respect to a change in the mass of the solute is then characterized by the partial specific volume (PSV) *via*

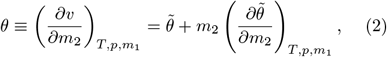

as motivated in [24].

For the sake of simplicity, for all the following considerations, we employ the concept of volume additivity, for which it is straightforward to show that 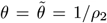, where *ρ*_2_ is the solute density (see SI; Eqs. (S3) [25]). By denoting the solute concentration as *c*_2_ ≡ *m*_2_*/v* and expressing the mass of the solvent as

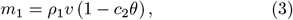

we obtain an expression for the MD of the mixture as

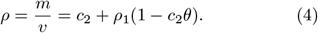

In the next step, we connect the RI of the solution to the solute concentration *c*_2_ *via* the phenomenological mixing rule

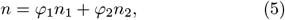

with *φ*_*i*_ = *c*_*i*_*/ρ*_*i*_ = *v*_*i*_*/v* being the volume fraction of component *i* [26]. In the following, we refer to Eq. (5) as Biot equation or mixing rule. Note that in a typical ODT experiment we determine the RI contrast *δn*≡ *n* −*n*_1_ which has to fulfill a non-negativity constraint (i.e., *δn*≥ 0). Evaluating Eq. (5) while assuming volume additivity [(1 − *φ*_2_) = *φ*_1_], we arrive at

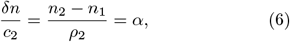

where we identify the customarily designated RI increment *α*. Finally, by replacing *c*_2_ in Eq. (4) with the expression given in Eq. (6), we obtain an estimate of the solution MD in dependence of the RI as

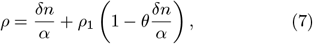

which has been employed in [17], or stated equivalently in [27, 28]. While relations similar to Eq. (7) can be found for different RI mixing rules (see e.g., [29]), the assumption of volume additivity is central for our considerations. Eq. (7) is the basis of all following considerations and will be employed frequently throughout this study.

The first term of Eqs. (7) and (4), respectively, computes the solute concentration and the second term accounts for the MD of the solvent as well as the volume uptake of the solute content. In other words, the second term can be interpreted as the MD contribution of the solvent content of the sample. In this interpretation, the PSV, *θ*, scales the solvent content.

Experimentally, the PSV of a solute is determined by measuring the density of the solution *ρ* in dependence of the solute concentration *c*_2_ [24], which can be motivated by solving Eq. (4) for *θ* as

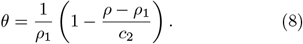

Hence, any experimentally observed deviation from a constant slope of *ρ*(*c*_2_) can be directly interpreted as a violation of the volume additivity assumption.

A straightforward and common approach to obtain the MD of a liquid is given by pycnometry, where the mass of a precisely fixed volume is measured. The MD is then simply given by the mass to volume ratio. Further methods to measure MD are reviewed in [24] and references therein. Similarly, the RI increment is customarily determined by measuring the RI of the solution in dependence of the solute concentration (see Eq. (6)), e.g., by employing an Abbe refractometer.

In the context of biological matter, one may think of the ‘solute’ as a complex composition of many constituents, which makes the experimental determination of both, PSV and RI increment for all components and their combinations practically impossible; the human proteome alone consists of ∼ 10^4^ proteins [18, 30]. In order to resolve this problem, at least partially, not accounting for volume inadditivities, a method of determining the correlative distributions 𝒫 (*α, θ*) of the proteomes of different organisms was introduced and experimentally validated for two amino acid sequences in [23]. The authors computed the weight averages of the residue refractivity per gram and according PSV *θ* of proteins based on their respective amino acid sequence. The refractivity per gram *R*_*i*_ of a molecule *i* is proportional to its polarizability 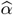 and molar mass *M* as 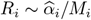. Further, the refractivity per gram can be connected to the RI and the MD of the solute *via* the Lorentz-Lorenz (or Clausius–Mossotti) relation (see e.g., [31]) as

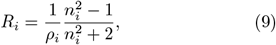

which may be solved for the RI of the molecule *n*_*i*_, to obtain

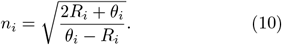

Considering that the PSV of a binary solution is given by the mass average of the PSVs of the solvent and solute as

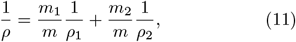

consequently, one could assume that the refraction per gram of the solution is given by the mass average of the constituents as well

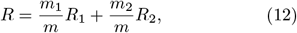

as it carries the same unit [ml/g] [32]. By inserting Eq. (9) into Eq. (12), we obtain the customarily denoted Lorentz-Lorenz mixing rule of RIs

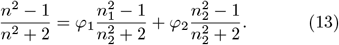

We note that Eq. (13) could be employed to derive an expression of the MD in dependence of the RI, similar to Eq. (7), using the Biot Eq. (5). Furthermore, we want to stress that neither the Lorentz-Lorenz mixing rule of RIs (Eq. (13)), nor the Biot mixing rule of RIs (Eq. (5)) do explicitly require volume additivity (see [26, 33] for further details).

However, assuming dilute solutions (*n* → *n*_1_) as well as volume additivity, the authors of [23] computed the RI increment for each protein, denoted by the index p, as derived in [26], as

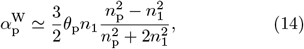

by employing the Wiener mixing rule of RIs

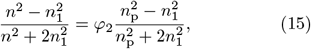

and Eq. (10) to compute the RI of each protein *n*_p_ from the mass averaged refractions per gram and PSVs of the respective amino acid sequences, as described earlier. Repeating this procedure for all proteins presumed to be abundant in the different organisms under study, they obtained the bivariate distribution of RI increment and PSV. Zhao *et al*. [23] then fitted normal distributions to the univariate histograms to obtain the means and standard deviations for different organisms, of which we depicted two in Tab. I. We repeated the computations presented in [23] (see SI [25]) for the updated human and zebrafish proteomes obtained from [30]. Besides computing the RI increment from the Wiener relation for dilute solutions (Eq. (14)), we also employed the Biot Eq. (5) in combination with the volume additivity assumption to obtain an expression of the RI increment as given in Eq. (6)

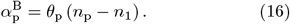

Furthermore, different to [23], we employed the consensus averages for the amino acid residue molecular volumes of [34]. The full list of parameters employed here is given in Tab. SII [25]. The resulting bivariate distributions 𝒫 (*α, θ*) are given in Fig. 1. The corresponding mean values and standard deviations of the fits of the univariate histograms with a normal distribution are given in Tab. I. Evidently, the PSV and RI increment values obtained here coincide very well, with the values from [23]. However, since our values were computed using the consensus values of the molar residue volumes of the amino acids of [34] instead of the ones of Cohn and Edsall [35], they are systematically lower. Furthermore, the RI increments obtained using Eq. (16) are systematically higher than the ones obtained from the Wiener equation for dilute solutions, given in Eq. (14).

**FIG. 1:**
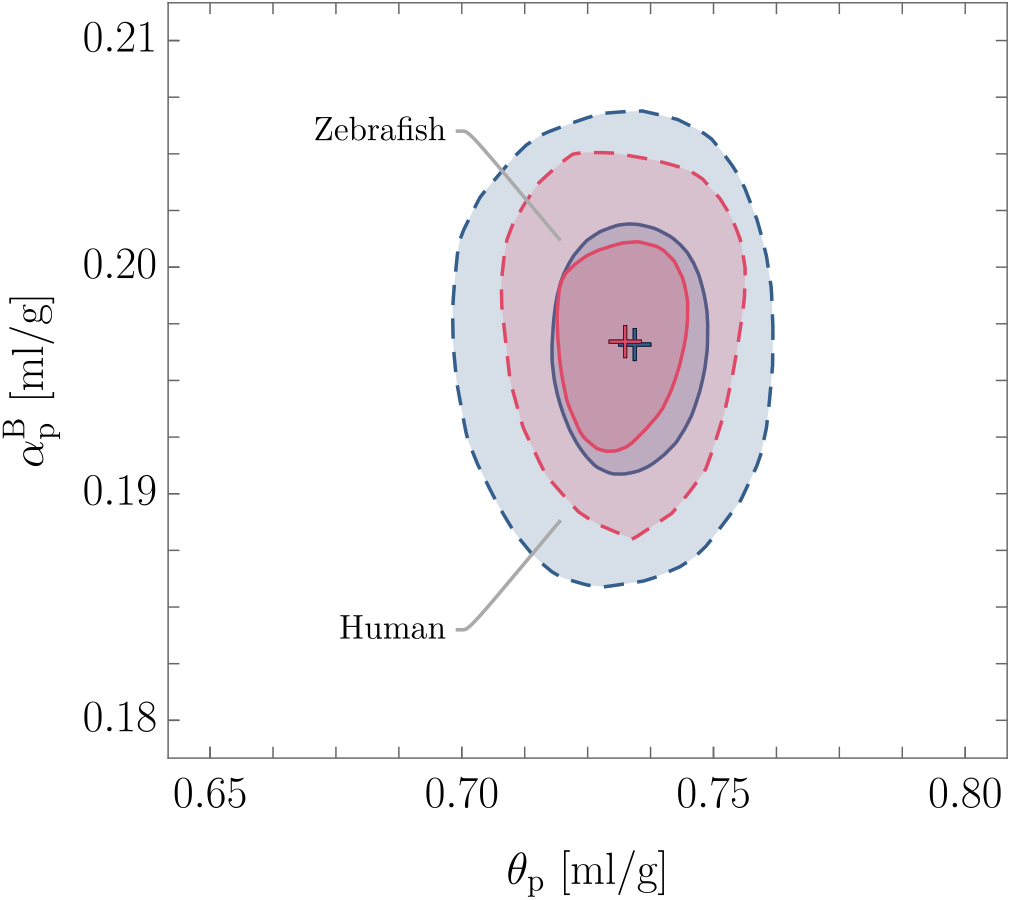
1*σ* (solid) and 2*σ* (dashed) likelihood contours of the correlated distributions of RI increment 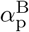 in ml/g and PSV *θ*_p_ in ml/g, for the human proteome; 82127 proteins (blue), and the zebrafish proteome; 46517 proteins (red).

While it is not clear why the calculations proposed in Zhao *et al*. [23] result in distributions 𝒫 (*α, θ*) that resemble uncorrelated bivariate normal distributions for the proteomes of different species, they facilitate the idea of taking the whole distribution 𝒫 (*α, θ*) into account when estimating the MD *via* Eq. (7). Consistently, in order to obtain a reliable estimate of the MD distribution, a precise characterization of the solute composition is needed.

To our knowledge, the justification for or against the assumption of volume additivity in complex biological matter is yet to be given (experimentally). However, we may interpret Eq. (2) in first order approximation as

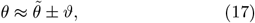

where any potential deviation from the volume additivity *ϑ* could be treated as an additional factor that will also contribute to the uncertainty of the RI and MD estimates.

In the following, we construct and employ a theoretical framework in which the information of the solute composition is incorporated into the prediction of the MD. For that purpose, we employ the Biot Eq. (5) for the majority of our further considerations and denote *α* = *α*^B^.

### III. EXTENSION OF THE BINARY MIXTURE MODEL

As motivated earlier, when dealing with biological matter, the complexity of the solute composition should be taken into account in order to obtain reliable estimates of the MD. To that aim, we first extend the expression of the MD of the binary mixture (Eq. (7)) to the case of different solute constituents, e.g., proteins of the human proteome, lipids and sugars, being dissolved in a solvent with corresponding RI *n*_1_ and MD *ρ*_1_.

We describe this problem by discretizing the total sample volume into *N*_v_ voxels with volumes *v*_v_

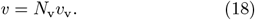

Furthermore, we discretize the voxels into *N*_0_ ‘voxelinos’ with volumes *v*_0_ as

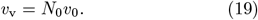

The motivation to divide a voxel into *N*_0_ voxelinos is to obtain small, yet macroscopic, volume fractions with a constant volume *v*_0_ that contain one and only one solution constituent. Hence, each voxelino can be either a solvent voxelino, or a solute voxelino and is inherently characterized by its respective partial specific volume *θ*_*i*_ and refraction per gram *R*_*i*_, i.e., its MD and RI. The number of solvent voxelinos in the voxel, *N*_1_, is given by *N*_1_ = *N*_0_− *N*_s_, where *N*_s_ is the number of solute voxelinos. Accordingly, the solvent volume fraction of a voxel is given by

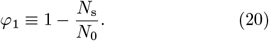

Accounting for multiple types of solute molecules, e.g., proteins, lipids, sugars, etc., being present in the solution, we choose the PSVs and refractions per gram of the solute voxelinos *θ*_*i*+1_ and *R*_*i*+1_ to be random values, drawn from a weighted mixture distribution

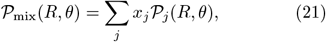

where *j* denotes the solute constituents. Further, the 𝒫_*j*_(*R, θ*) represent the bivariate probability distributions of the refraction per gram and PSV of the respective constituents and the associated weights *x*_*j*_ are given by the relative volume fractions as

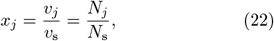

where *v*_s_ denotes the total solute volume. We note that Eq. (21) should be seen as a way of denoting that *N*_*j*_ out of *N*_s_ solute voxelinos of constituent *j* are present in a voxel. Consequently, *N*_*j*_ solute voxelinos have the RI *n*_*j*_ and MD *ρ*_*j*_ and we have that *N*_s_ = Σ_*j*_ *N*_*j*_, i.e., Σ_*j*_ *x*_*j*_ = 1.

Employing Eqs. (4) and (5), we readily obtain the relation between the MD *ρ* and the RI *n* of one voxel

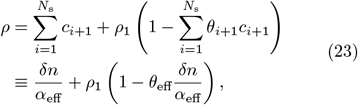

where we defined the effective RI increment *α*_eff_ and PSV *θ*_eff_. The RI of the solution *n* is then given by the Biot Eq. (5), where the RIs of the individual solute voxelinos *n*_*i*+1_ could be known directly from RI measurements, or may be computed by employing the Lorentz-Lorenz relation, given in Eq. (10).

It can be shown (see SI; Eqs. (S17) and (S18) [25]) that the effective parameters may be expressed by the mass averages of the respective parameters of the solute voxelinos as

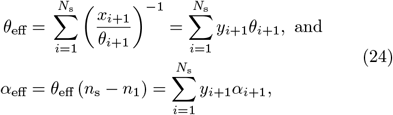

where we denoted the solute concentration and RI by *c*_s_ and *n*_s_, respectively, and the relative mass fraction of a solute voxelino by

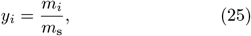

with *m*_s_ being the total solute mass. Hence, the effective parameters absorb the ‘complexity’ of the mixture, while the functional relationship of Eq. (4) is obeyed. With this at our disposal, we are able to compute the MD *ρ* and the corresponding RI *n* of a voxel, given a distribution 𝒫_mix_(*R, θ*) and a solvent volume fraction of the voxels *φ*_1_, i.e., the ratio *N*_s_*/N*_0_. Repeating this procedure for *N*_v_ voxels provides a map of RI values in resemblance of an ODT measurement, alongside the corresponding MDs.

As the framework introduced above is strongly dependent on *a priori* assumptions of the individual model parameters (*φ*_1_, *x*_*j*_, *α*_*j*_, *θ*_*j*_) for different complex mixtures, in the next step, we shall discuss the impact of these assumptions on the uncertainties of the RI and MD predictions.

## IV UNCERTAINTY QUANTIFICATION OF THE EXTENDED MODEL

Given access to experimental data, i.e., ODT tomograms of a sample of interest, we may not only compare the mean values of the measured and predicted RI distributions, but also their widths. This in turn enables a more reliable estimate of the MD distribution. Hence, in the next step we study the dependence of the uncertainty of the MD estimate *ρ*, defined in Eq. (23), and the RI *n* on the sample properties, namely, the solvent volume fraction *φ*_1_, the effective RI increment *α*_eff_, the effective partial specific volume *θ*_eff_ as well as their respective associated uncertainties.

To that aim, we investigate the impact of the presence of a second type of macro molecule in the mixture of proteins and water. Since lipids make up for about 13% of the solute mass fraction in mammalian cells [18], and typically exhibit an MD lower than water, they merit detailed scrutiny. In the following, we assume that the lipids are present in the form of lipid droplets and form an emulsion in the water+protein phase. For the sake of simplicity, we further assume that the lipid droplets consist only of the neutral lipid triolein (TO), neglecting sterol esters, triacylglycerols and phospholipids [36– 38]. The corresponding values of the RI, refraction per gram, PSV and RI increment of the two types of solute molecules under investigation are given in Tab. SI [25].

Throughout this study we assume that both RI and MD of the solvent are precisely known. We further point out that we implement the values given in Tab. SI [25] in our calculations as follows: if a quantity is stated as mean *±* standard deviation, we account for it as normally distributed with respective mean and standard deviation. For the cases where we could not estimate an uncertainty, we assume the quantity to be delta distributed.

### A. Effective RI increment and PSV

Examining the definitions of *α*_eff_ and *θ*_eff_, given in Eqs. (24), we observe a dependence of both quantities on the number of solute voxelinos per voxel *N*_s_, given as the upper limit of the sum. To obtain an intuition about the implications on the respective uncertainties, we first consider the case of proteins dissolved in water, which may be approximated by uncorrelated normal distributions 𝒩 (*μ, σ*) of the RI increment and PSV, as shown in Fig. 1

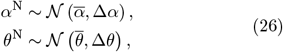

with respective mean values and standard deviations (see e.g., Tab. I). Applying GPU to Eqs. (24) we have

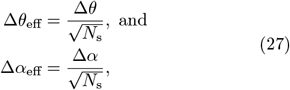

where we used that the relative mass fraction of each solute voxelino is given by *y*_*i*+1_ = 1*/N*_s_. The result of Eq. (27) is in concordance with the central limit theorem (CLT). In other words, the standard deviation of the effective distributions corresponds to the ‘standard error of the mean’ of the initial distributions, given that the voxel contains *N*_s_ protein voxelinos. Thus, considering that in biological matter the number of voxelinos per voxel is typically larger than ∼ 10^8^ for experimentally accessible voxel sizes in the order of 1 μm^3^, the deviations of the effective RI increment and PSV, for the case of proteins dissolved in water, are negligibly small.

Next, we study the impact of lipids in the protein+water mixture. In this scenario, Eq. (21) takes the form

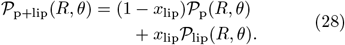

Assuming that *𝒫*_p_ and *𝒫*_lip_ follow uncorrelated bivariate normal distributions, using Eqs. (27), we find that the distributions of the effective PSV and RI increment follow normal distributions as

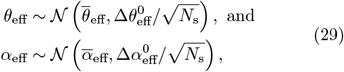

where the respective mean values of effective PSV and RI increment are given in Eqs. (24) and the standard deviations 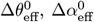 follow the standard deviation of a mixture distribution, provided in the supplemental information (Eq. (S14) [25]).

With this, we further examine the implications of deviations of the relative lipid volume fraction ∆*x*_lip_ from voxel to voxel. Such deviations may be interpreted as a form of inhomogeneity of the system, which have been experimentally quantified in cells and tissues by SRS measurements [20]. Employing GPU we find the following analytical expression of the deviation of the mean effective PSV and RI increment with respect to ∆*x*_lip_ as

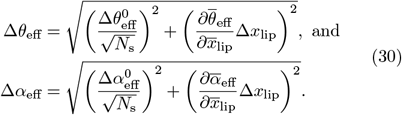

The partial derivatives in Eqs. (30) are given in the supplemental information (Eqs. (S19) and (S20) [25]). A visual depiction of Eqs. (30) in dependence of the number of voxelinos per voxel *N*_0_ = *N*_s_*/*(1− *φ*_1_) and the corresponding results of MC simulations for certain parameter configurations is shown in Fig. 2(a).

**FIG. 2:**
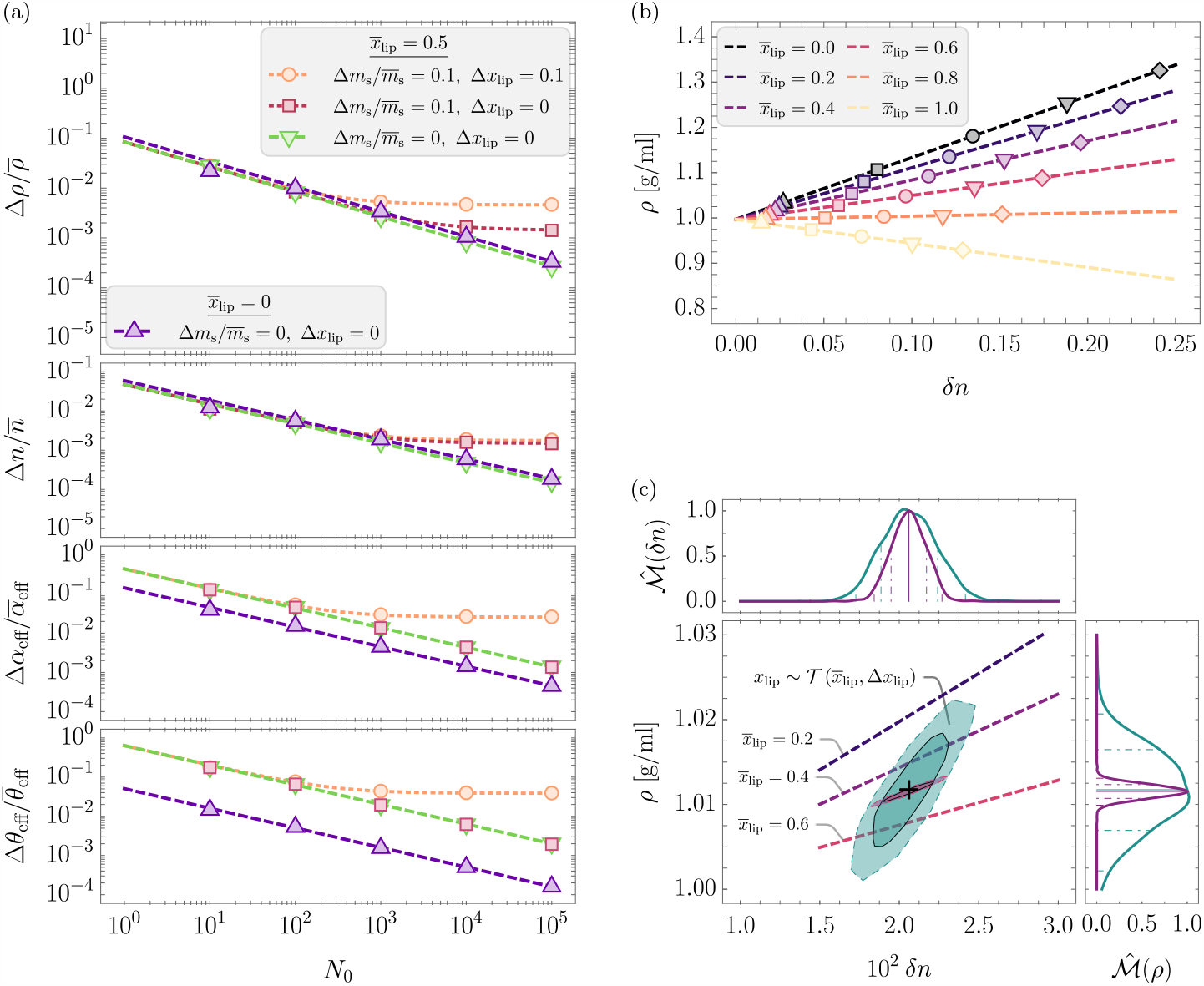
Results of MC simulations and according analytical solutions for the mixture of human proteins and the neutral lipid triolein in water; (a): Relative deviations of the MD *ρ*, the RI *n*, the effective RI increment *α*_eff_ and PSV *θ*_eff_ in dependence of the number of molecules per voxel *N*_0_ obtained from MC simulations (symbols) and analytical solutions (dashed lines) for different mean relative lipid volume fractions 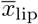, associated deviations ∆*x*_lip_ and relative deviations of the number of the solute mass 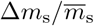. The MC simulations were performed for a mean water volume fraction of 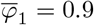 and *N*_v_ = 10^3^. (b): MD *ρ* in g/ml in dependence of the RI contrast *δn* for different mean relative lipid volume fractions *x*_lip_ and mean water volume fractions 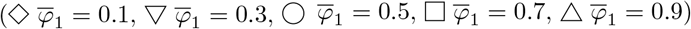 for ∆*x*_lip_ = 0, *N*_0_ = 10^3^ and *N*_v_ = 10^2^. The dashed lines indicate the analytical solutions of Eq. (23). (c): Correlative distribution of the MD *ρ* in g/ml and the RI contrast *δn* (the solid and dashed lines indicate the 68.3% and 95.5% confidence contours) with the corresponding normalized marginal probability density distributions 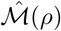 and 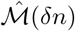, respectively (the solid line represents the median, the dash-dotted and dashed lines indicate the 68.3% and 95.5% confidence intervals), for the exact relative lipid volume fraction 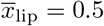 (purple) and the relative lipid volume fraction following a truncated normal distribution 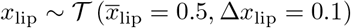 (cyan, see main text). The MC simulations were performed for a water volume fraction following a truncated normal distribution 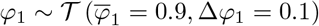, *N*_0_ = 10^5^ and *N*_v_ = 10^3^.

As becomes apparent, the analytical solution, employing GPU, is in concordance with the simulated values. However, we note that this is due to the assumption of normal distributions for the individual effective RI increments and PSVs. For non-normal distributions, GPU might not be applicable.

By assuming the mixture distribution, given by Eq. (28), consequently, the deviations ∆*α*_eff_ and ∆*θ*_eff_ are maximized for 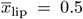 (see SI [25]). Secondly, while for non-fluctuating *x*_lip_, i.e., ∆*x*_lip_ = 0, the respective deviations of the effective RI increment and the PSV exhibit the 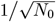 proportionality, for ∆*x*_lip_ *>* 0, we obtain non-vanishing deviations for a large number of voxelinos per voxel *N*_0_.

Throughout this study we assume that all volume fractions, in particular *x*_lip_, follow a normal distribution, truncated in the domain [0, 1], since values outside of this interval are non-physical under the assumption of volume additivity. For a normal distribution with mean *μ* and standard deviation *σ*, the corresponding truncated distribution is denoted by *𝒯* (*μ, σ*). The probability density function (PDF) of 𝒯 is defined as

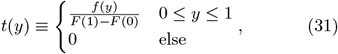

where *f* (*y*) and *F* (*y*) denote the PDF and cumulative distribution function (CDF), respectively, of said normal distribution 𝒩 (*μ, σ*). We note that another choice of distribution could be given by the beta distribution ℬ (*a, b*) with shape parameters *a* and *b*, which is inherently bounded between 0 and 1. However, the interpretation of the shape parameters in this context is not as straightforward as the truncated normal distribution. Alternatively, a uniform distribution with a certain domain 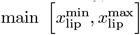 could be employed. For more than two types of macro molecules, the Dirichlet distribution could be of use. However, we expect a similar qualitative behaviour for all mentioned distributions.

### B. Solvent volume fraction

Next, we study the uncertainty associated to the solvent volume fraction *φ*_1_, defined in Eq. (20), in dependence of the number of solute voxelinos *N*_s_ per voxel. By employing GPU we find

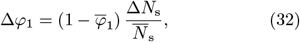

where we used the presumption that the number of voxelinos per voxel *N*_0_ is constant. From Eq. (32) we obtain that a change in the solvent content from voxel to voxel is directly proportional to a relative cha<u>n</u>ge in the number of solute voxelinos per voxel 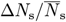. However, it is not clear whether *N*_s_ follows a statistical distribution, and if so, how this distribution would be governed by active/passive processes in biological matter. An ingenuous guess is given by the equilibrium assumption that the number of solute voxelinos per voxel *N*_s_ is binomially distributed as

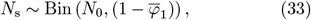

which may be interpreted as finding *N*_s_ out of *N*_0_ voxelinos in a voxel with a probability of 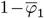. The according standard deviation and mean value of *N*_s_ is then given by

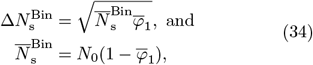

respectively, which is consistent with Eq. (20). Besides a statistical argument, we may also compute a change in the number of solute voxelinos from voxel to voxel by considering

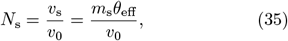

from which we have

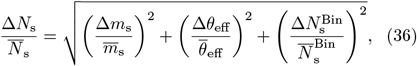

using the presumption of a constant voxelino volume ∆*v*_0_ = 0. Herewith, Eq. (32) may be written as

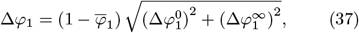

with

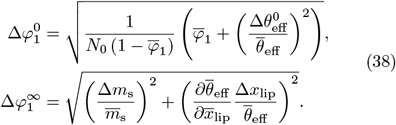

This indicates that, similarly to the effective RI increment and PSV, given by Eqs. (27), the deviation of the solvent volume fraction ∆*φ*_1_ h<u>as t</u>wo components; for once 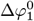, which shows the 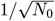 dependence, following the CLT. Secondly, 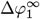, which connects fluctuations in the solvent volume fraction to fluctuations in the solute mass and/or fluctuations in the solute composition. Hence 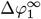 may be interpreted as quantification of the degree of inhomogeneity of the sample in the solute and its composition. Consequently, for a large number of voxelinos per voxel *N*_0_, these inhomogeneities become the dominant contribution to the deviation of the solvent volume fraction.

We note that the presumption of a constant voxelino volume ∆*v*_0_ = 0 and a constant number of voxelinos per voxel ∆*N*_0_ = 0 from voxel to voxel is necessitated by the experimental boundary condition, that all voxels have the same volume, i.e., ∆*v*_v_ = 0.

### C. Refractive index

Considering the previous derivations of the uncertainties of the effective PSV and RI increment, as well as the water volume fraction, we now examine the refractive index of the solution. For that purpose, we rewrite Eq. (5) as

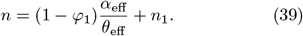

With this at hand, we readily obtain an estimate of the uncertainty of *n* by employing GPU (neglecting potential correlations) as

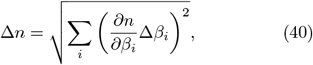

where the sum is taken over all *β* = {*α*_eff_, *θ*_eff_, *φ*_1_} and the respective deviations ∆*β*_*i*_ are given in Eqs. (30) and (37). The graphical representation of Eq. (40) in dependence of *N*_0_ is given in Fig. 2(a).

As a consequence we have that for a vanishing relative deviation of the solute 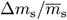, the 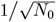 scaling behaviour of the effective RI increment and PSV determines the scaling behaviour of the deviations in the RI ∆*n*. However for 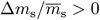 we obtain a constant deviation of the RI for large *N*_0_, consistent with the broad RI distributions obtained by ODT measurements of different cells and tissues [12, 15, 19, 39], and fluctuations in the water volume fraction measured by SRS [20]. Furthermore, while the dependence of ∆*n* on the mean relative lipid volume fraction 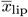 is determined by the deviations of the effective RI increment and PSV, we observe a vanishing impact of the deviation of the relative lipid volume fraction ∆*x*_lip_. This fact is due to the numerically small differences of the mean refractions per gram of the particular choice of proteins and the lipid 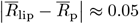 in combination with larger differences of the mean PSVs 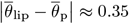 (see SI; Fig. S4 [25]).

### D. Mass density

Finally, we investigate the uncertainty associated with the MD. For convenience, we may express Eq. (23) as

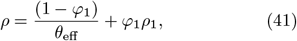

to obtain the corresponding deviation, employing GPU, as

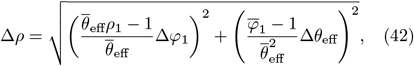

displayed in Fig. 2(a). As discussed for ∆*n*, the magnitude of ∆*ρ* is impacted by the mean relative lipid fraction 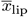 as a consequence of the mixture distribution, given in Eq. (21). Furthermore, we have a non-vanishing deviation of the MD for 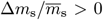 and large *N*_0_. To no surprise, we observe a remarkable impact of deviation of the relative lipid fraction ∆*x*_lip_ on ∆*ρ* due to the strong scaling with the deviation of the effective PSV ∆*θ*_eff_.

### E. Correlation of refractive index and mass density

Having studied the deviations associated with RI and MD, we next sought to illuminate the correlation between the distributions of the RI contrast *δn*, and the MD *ρ*, denoted by 𝒫 (*ρ, δn*), in dependence of the model parameters introduced earlier. To that aim, we performed MC simulations of Eq. (23) for the case of human proteins and triolein in water for a range of different mean relative lipid and water volume fractions, as shown in Fig. 2(b).

Besides the intuitive behaviour of *ρ*(*δn*) for the cases of 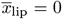 (MD increases with increasing refractive index) and 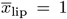 (MD decreases with increasing refractive index), for 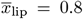 the MD is roughly constant for all refractive index values. This feature is quite remarkable since it demonstrates that for certain solute compositions, the MD is decoupled from the refractive index for all possible water volume fractions. Furthermore, as motivated earlier, and shown in Fig. 2(c), measuring a refractive index distribution, e.g., *via* ODT, may correspond to a range of different water and relative lipid volume fractions, resulting in drastically different estimates on the distribution of the MD from case to case. This in turn strongly motivates the necessity for detailed knowledge about not only the solute composition, but also the solvent content of the sample.

In the light of the theoretical implications delineated above, in the next step we want to examine the predictive capabilities of the model for a set of physiological complex mixtures that are well characterized in terms of their solute composition.

## V. EXPERIMENTAL VALIDATION AND APPLICATION

In the following, we scrutinize the applicability of previous findings to a set of well characterized, physiological substances, i.e., *bovine* skim-milk powder (SM, Millipore 70166) and 20% intralipid emulsion (IL, Sigma-Aldrich I141). To that aim, we measured the solute concentration-dependent RI and MD with an Abbe refractometer and a pycnometer, respectively. Both samples are particularly intriguing since they should exhibit different *ρ*(*δn*) dependencies; SM mainly consists of lactose and milk proteins, while IL is a stabilized emulsion of soybean oil (see Fig. 2(b) for a reference).

According to the chemical certificate of analysis, provided by the manufacturers, the SM exhibits 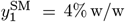 and the IL exhibits 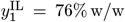of water.

Hence, we computed the respective solute concentrations as

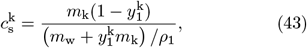

where *m*_k_ denotes the mass of the sample (SM or IL) and *m*_w_ is the mass of water added to the sample. The results of the measurements are shown in Figs. 3(a) and 3(b), where each point represents *N* = 5 technical repetitions. Using Eqs. (4) and (6) we fitted the data *via* a *χ*^2^-minimization approach to obtain experimental values of *θ*_eff_ and *α*_eff_ with according uncertainties, respectively, for both, SM, and IL. Examining the fitting residuals, evidently, the RI contrast and MD scale linearly with the solute concentration, justifying the assumption of volume additivity.

**FIG. 3:**
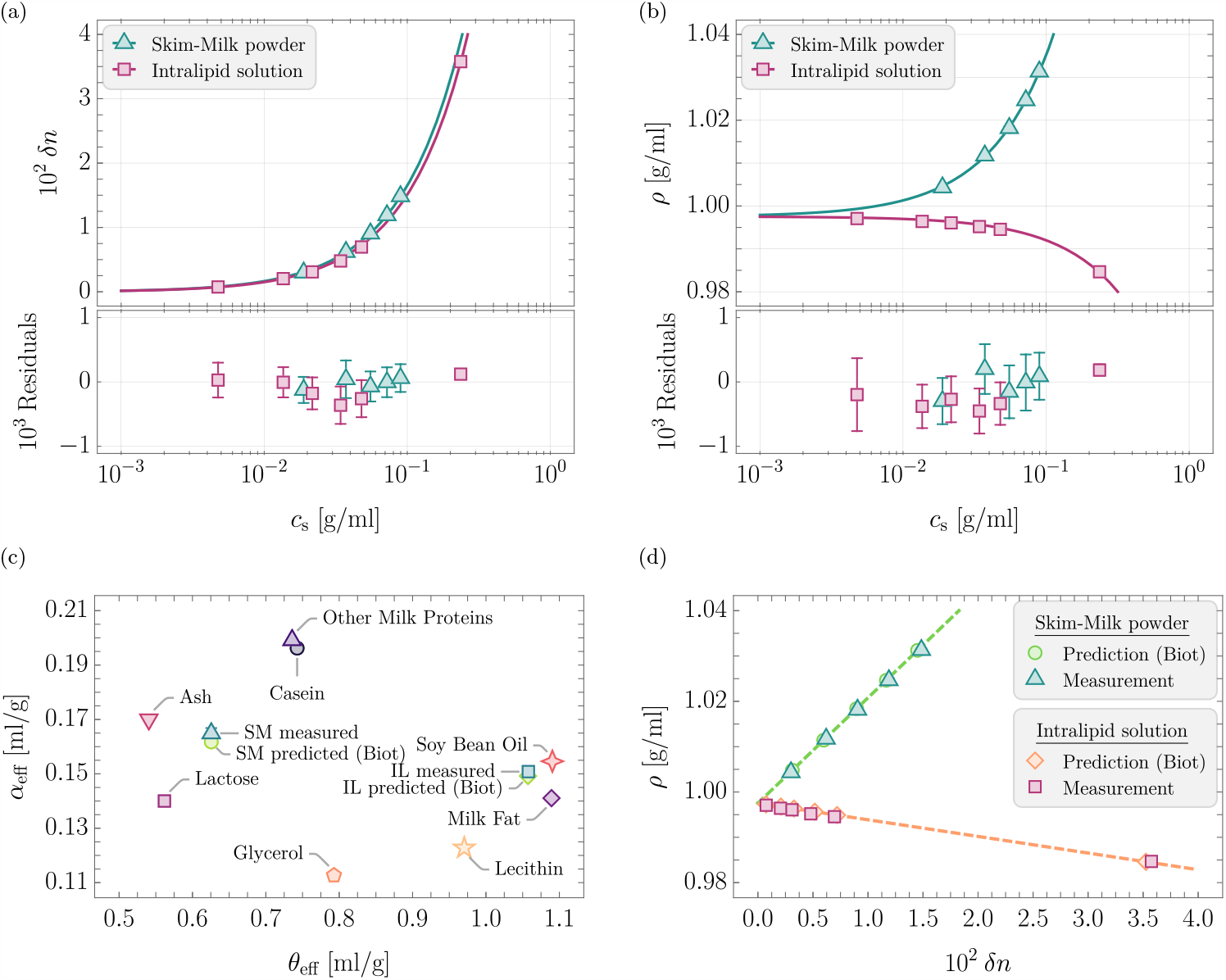
Results of the concentration dependent measurements of the MD and RI of *bovine* skim-milk powder (SM) and 20% intralipid emulsion (IL) in water as well as the theoretical predictions based on the chemical composition using Eq. (23); (a): RI contrast *δn* in dependence of the solute concentration *c*_s_ in g/ml of SM (Δ, *N* = 5 technical repetitions) and IL (◻, *N* = 5 technical repetitions) with the respective fits of Eq. (16) (solid lines) and fit residuals. (b): MD *ρ* in g/ml in dependence of the solute concentration *c*_s_ in g/ml of SM (Δ, *N* = 5 technical repetitions) and IL (◻, *N* = 5 technical repetitions) with the respective fits of Eq. (4) (solid lines) and fit residuals. (c): Effective RI increment *α*_eff_ in ml/g and PSV *θ*_eff_ in ml/g of various substances that compose SM and IL, as well as the measured and predicted values for SM and IL. (d): MD *ρ* in g/ml in dependence of the RI contrast *δn* for different concentrations of SM in water and IL. The symbols represent measured values (*N* = 5 technical repetitions) and the predicted values using the Biot mixing rule (Eq. (5)) for *N*_0_ = 10^4^ and *N*_v_ = 10^3^. The dashed lines indicate Eq. (23) for the predicted values of the effective RI increment and PSV.

We then employed the information about the chemical composition of the respective substances, provided by the manufacturers (see Tab. SI [25]), to compute the correlative RI contrast and MD according to Eq. (23) and the Biot RI mixing rule (Eq. (5)), employing MC sampling. A graphical representation of the respective RI increments and PSVs of all substances considered, as well as the experimentally determined and predicted values of SM and IL is given in Fig. 3(c). The numerical values and respective references are stated in Tab. SI [25]. As becomes apparent, the measured and predicted values of the SM and IL are in good agreement, while potential uncertainties regarding the exact chemical composition might be underappreciated here as we have no means of estimating them.

Furthermore, matching the experimental concentrations, we obtained a prediction of the MD in dependence of the RI, which was found to coincide well with the measurements for both SM, and IL, as shown in Fig. 3(d).

### A. Larval zebrafish trunk tissue

Having delineated the good agreement between the assumptions used to derive an expression of the MD in dependence of the RI for two dilute, physiological samples, we now want to examine the capability of the proposed model in an *in vivo* scenario. To that aim we chose the larval zebrafish model system at 96 hours postfertilization (hpf), for which mass spectrometry (MS) and RI data, employing ODT, of the trunk tissue were recently obtained [19]. The RI data is given in Tab. SIII [25]. As the MS data provides the protein content, we are able to estimate the RI increment and PSV distribution of the proteins present in the tissue, as demonstrated earlier, following Zhao *et al*. [23] (see Tab. I).

In the following, we assume that the larval zebrafish trunk tissue primarily consists of mentioned proteins, lipids and water, based on the findings of [40, 41]. The lipid composition of zebrafish larvae was determined in [41], from which we adapted the four phospholipid fatty acids (PLFA) with the highest abundance. The respective relative volume fractions of the PLFAs for zebrafish larvae at 96 hpf were roughly digitally obtained from [41] and are provided in Tab. SI [25]. These four PLFAs make up for about 72% of the total PLFA amount in the zebrafish larva. Additionally, based on [41], we assume that the overall lipid composition of the tissue is only given by triolein (TO) and said PLFAs.

Next, we inferred the distributions of the relative lipid volume fraction *x*_lip_ and the water volume fraction *φ*_1_ based on the measurements of [40] (see SI [25]). From these initial estimations, employing Eq. (23), we obtained the correlative RI and MD distribution, displayed in Fig. 4(b), by MC sampling.

**FIG. 4:**
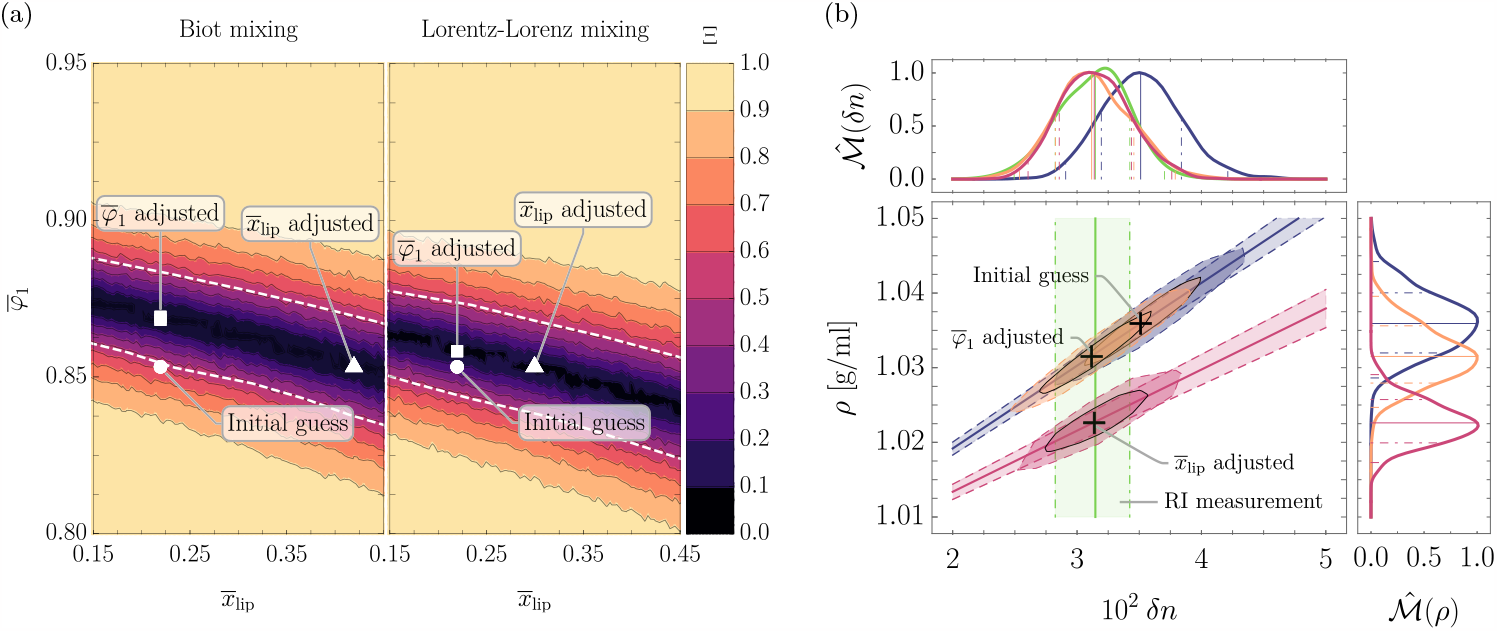
Results of MC simulations and according analytical solutions for the trunk tissue of larval zebrafish; (a): Quantile comparison effect size (QCES) Ξ between the RI distribution of larval zebrafish trunk tissue measured in [19] and the predictive RI distributions obtained from MC simulations of Eq. (23) using the Biot mixing rule (left, Eq. (5)) and the Lorentz-Lorenz mixing rule (right, Eq. (13)) for different mean water volume fractions 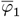 and mean relative lipid volume fraction 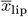. The symbols indicate ◯: the initial guess, based on the measurements of [40], ◻: the local minimum of Ξ for the relative lipid volume fraction of the initial guess and △: the local minimum of Ξ for the water volume fraction of the initial guess. The white dashed lines indicate a 1 *σ* deviation of the predicted RI distribution to the measured RI distribution. All MC simulations were performed for *N*_0_ = 10^3^ and *N*_v_ = 10^3^. (b): Predicted correlative distributions of the MD *ρ* in g/ml and the RI contrast *δn* of larval zebrafish trunk tissue with the corresponding normalized marginal probability density distributions 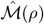and 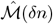, respectively (the solid line represents the median, the dash-dotted and dashed lines indicate the 68.3% and 95.5% confidence intervals). The ellipsoids indicate the different cases shown in Fig. (a): the initial guess (blue), 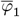 adjusted (orange) and 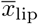 adjusted (red) with the corresponding analytical predictions for the different relative lipid volume fractions and varying water volume fraction. The green band indicates the RI measurement of [19]. The MC simulations were performed for *N*_0_ = 10^5^ and *N*_v_ = 10^3^.

**Table 1:** Mean and standard deviations of the RI increments 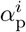 and partial specific volumes *θ*_p_ distributions based on amino acid sequences of the proteome of the human and the zebrafish and the trunk tissue of the larval zebrafish at 96 hpf. The calculation of 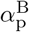 employed the Biot equation given in Eq. (5), while 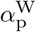 was derived from the dilute limit of the Wiener mixing rule of refractive indices (Eq. (14)).

| | $\bar{\alpha}_p^W$ in ml/g | $\Delta\alpha_p^W$ in ml/g | $\bar{\alpha}_p^B$ in ml/g | $\Delta\alpha_p^B$ in ml/g | $\bar{\theta}_p$ in ml/g | $\Delta\theta_p$ in ml/g |
| --- | --- | --- | --- | --- | --- | --- |
| Human |  |  |  |  |  |  |
| Proteome<sup>a</sup> | 0.1899 | 0.0030 | N/A | N/A | 0.735 | 0.010 |
| Proteome<sup>b</sup> | 0.188 | 0.004 | 0.196 | 0.004 | 0.734 | 0.012 |
| Zebrafish |  |  |  |  |  |  |
| Proteome<sup>a</sup> | 0.1904 | 0.0030 | N/A | N/A | 0.735 | 0.010 |
| Proteome<sup>b</sup> | 0.1887 | 0.0031 | 0.1968 | 0.0034 | 0.732 | 0.010 |
| trunk tissue<sup>b</sup> | 0.1883 | 0.0029 | 0.1963 | 0.0032 | 0.734 | 0.009 |

<sup>a</sup> [23]
<sup>b</sup> this work

Evidently, both measured and predicted RI distributions do not coincide within one standard deviation. This fact is not unexpected since the measurements of [40] were done on whole animals, including the yolk sac which is rich in lipids. Hence, the assumed relative volume fraction of the lipids and the water volume fraction are likely to be different from the trunk tissue. However, given the deviations on the distributions of the relative lipid volume fraction and the water volume fraction, the resulting deviation of the predicted RI distribution matches remarkably well with the deviation of the measured RI distribution.

Hence, in an attempt to find the mean water volume fraction 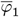 and the mean relative lipid volume fraction 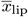 that resembles the experimental condition, we evaluated the quantile comparison effect size (QCES, see SI; Eq. (S11) [25]), as proposed in [42], denoted by Ξ, between predicted and measured RI distributions for a range of different 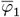 and 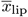, shown in Fig. 4(a). The QCES provides an elegant way of comparing any two distributions by means of the vertical quantile comparison divergence. It is bounded on the domain [0, 1]; for Ξ = 0 the two distributions are identical and for Ξ = 1, they are maximally distinct. To give further intuition on the QCES, we may consider two normal distributions with mean values *μ*_1_ and *μ*_2_ and the standard deviations *σ*_1_ = *σ*_2_ = *σ*. For the case of *μ*_2_ − *μ*_1_ = 1*σ*, which corresponds to a Cohen’s *d* = 1, the QCES Ξ ≈ 0.52. Hence, we consider any two distributions to be significantly different when Ξ *>* 0.52.

With this at our disposal, we obtain a region in the 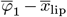 plane that establishes concordance between the predicted and measured RI distributions, as indicated by the white dashed lines in Fig. 4(a). Evidently, the MD is not well constrained by this set of parameters, since combinations of 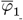 and 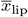 for a wide range of values result in the same RI, but drastically different MD distributions (see Fig. 4(b)). Consequently, minimizing the QCES under the assumption that either the initial guess of the water volume fraction or the initial guess of the relative lipid volume fraction is adequate, one obtains incompatible MD estimates, as indicated by the cases ‘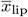 adjusted’ and ‘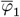 adjusted’ in Fig. 4(b). This fact convincingly shows the ambiguity of estimations of the MD given a measured RI distribution without correlative constraints on the (local) chemical composition of the sample. However, reconsidering that the measurements of [40] were performed on whole animals, including the yolk sac, the real relative lipid volume fraction of the trunk tissue should be equal or lower than the ‘initial guess’. This in turn makes the case of ‘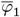 adjusted’ our best guess, yielding a MD estimate of *ρ*^B^ = (1.032 *±* 0.004) g/ml.

We note that, as indicated in Fig. 4(a), when employing the Lorentz-Lorenz mixing rule of RIs (Eq. (13)) instead of the Biot Eq. (5), the RI distribution obtained by the ‘initial guess’ is in good agreement with the measured RI distribution and we obtain an MD estimate for the case of ‘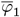 adjusted’ of *ρ*^LL^ = (1.034 *±* 0.004) g/ml, which is, considering the uncertainties, well compatible with the estimate we obtained employing the Biot Eq. (5). However, it is worth pointing out that the Lorentz-Lorenz mixing rule did not agree well with the RI measurements of the validation samples presented earlier.

In essence, at present, it is not clear which RI mixing rule should be employed in this *in vivo* scenario, without a more comprehensive understanding regarding the biochemical composition of the sample. However, once this insight becomes available, e.g., by measuring (S)RS, different mixing rules could be compared to each other, as presented earlier. This in turn would also allow to study whether the ‘best fitting’ RI mixing rule is conserved across different specimen.

Finally, we want to point out that if we use the customary simplifying assumption that the solute composition of the trunk tissue is only given by proteins, employing Eq. (7), the RI increment and PSV given in Tab. I as well as the measured RI data (Tab. SIII [25]), we obtain *ρ*_p_ = (1.040 *±* 0.004) g/ml, using GPU. Apparently, this value does not coincide well with the values determined above, *ρ*^B^ and *ρ*^LL^. In fact, as outlined earlier, not taking lipids into account results in a systematic overestimation of the mass density of biological matter.

## VI. DISCUSSION AND OUTLOOK

Considering the complexity of the chemical composition of biological matter in order to infer the ‘optical’ MD, based on a RI measurement is not well established in contemporary literature. Here we present a theoretical macroscopic model that is capable of describing the problem, employing a minimal set of assumptions, namely the Biot mixing rule of RIs and the assumption of volume additivity.

We evaluated the possible sources of uncertainties associated with the model and showed that based on the chemical composition of the sample and the associated degree of inhomogeneity, the resulting RI and MD distributions might drastically differ from the customary assumption of biological matter consisting of proteins and water only. For that purpose, we provided analytical solutions as well as consistent simulation results for the case of a binary solute, composed of proteins and lipids.

While it is shown that for singular proteins in solution the assumption of volume additivity might not be justified [43] (see SI [25]), we provided experimental evidence that, for the set of validation samples under investigation, i.e., *bovine* skim-milk powder and 20% intralipid emulsion, it holds well within the measurement uncertainties.

Further, we provided evidence that the predictions of the correlative MD and RI based on our model agree with the experimentally obtained values of the validation samples, thus establishing confidence in the application of the theoretical considerations presented here to estimate MD in dependence of the RI, given the chemical composition of the sample under study.

When applying the model to an *in vivo* specimen, i.e., the trunk tissue of the larval zebrafish, we observed that the mean value of RI measurements of [19] does not coincide within one standard deviation of our predictions, based on the estimations of the chemical composition of the tissue, employing the measurements of [40], where the authors determined the masses of water, proteins and lipids of whole animals. However, our initial predictions are remarkably close to the measurements, given the multitude of assumptions and simplifications employed.

We then explored the possibility of adjusting the hypothetical protein to lipid ratio and the water content of larval zebrafish trunk tissue in order to establish agreement between the measured and predicted RI distributions. To that aim we visited the topic of distribution comparisons. Employing the quantile comparison effect size, proposed in [42], we found a range of possible combinations of relative lipid volume fractions and water volume fractions for which the predicted RI distribution coincides with the measured one. However, for these inferred ranges, the MD varies significantly, resulting in large deviations. In order to resolve this problem, complementary (S)RS measurements would be necessary. With the information about the local concentrations of proteins and lipids in the trunk tissue, obtained by such (S)RS measurements, the predictive capability of our model would be significantly increased [20, 21].

Going forward, the estimation of the MD of biological matter from RI measurements, as outlined in this study, will have interesting implications for inferring the mechanical properties from opto-acoustical measurements, e.g., *via* Brillouin microscopy. The problem of a varying solute composition within a sample is well appreciated (see e.g., [12]), but not resolved in a cohesive manner. Evidently, combining Brillouin microscopy not only with ODT, but also incorporating the local biochemical composition (e.g., *via* (S)RS measurements) allows for a better estimation of the longitudinal (elastic) modulus. We note that the aforementioned problem could also be resolved by performing stimulated Brillouin microscopy in combination with ODT. Here, the MD can be obtained directly from measurements of the Brillouin resonance gain factor, which in turn is connected to the Brillouin gain and the pump laser power, as well as the RI, Brillouin frequency shift and Brillouin line width [44]. Accurately determining the local (*in vivo*) MD, will conceivably enable a more profound interpretation of functional mechanisms at play in biological matter.

The possibility of linking the MD estimates of a sample with the inherent dynamics of the system, i.e., the statistical processes, poses an intriguing endeavor. Moreover, explorations of the predictive RI and MD distributions of cells, in particular *Xenopus* egg extract, i.e., cytoplasm, for which the chemical composition could be accurately determined, will be insightful for inferences regarding the applicability and predictive capability of our model.

Finally, describing the problem of MD estimations, given uncertain *a priori* assumptions, in a Bayesian framework should be investigated.

In order to make the application of the findings of this study more accessible, we delineate strategies on how to estimate MD, given certain experimental paradigms, in the supplemental information [25].

## VII. MATERIALS AND METHODS

### A. Sample preparation

The skim-milk powder was dissolved in distilled water while carefully stirring the solution to avoid foaming. The solution was then left on a tilt/roller mixer for approximately 30 minutes. For the intralipid emulsion, according amounts of water were added to the emulsion and the solution was left on a tilt/roller mixer for approximately 30 minutes, as well. All samples were freshly prepared before the measurements were performed.

### B. Abbe refractometry and pycnometry

For measuring the refractive index of a liquid sample, 100 μl of the sample were loaded into an Abbe refractometer (KERN ORT 1RS) and a commercially available flashlight LED was employed as illumination source.

To determine the density of a liquid sample, a pycnometer (Blaubrand 43305) was employed. First, the volume of the pycnometer was determined by employing distilled water as a calibration sample (*N* = 10 technical repetitions) as

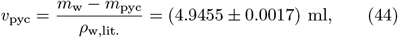

where *m*_w_ is the mass of the pycnometer filled with water, *m*_pyc_ is the mass of the empty pycnometer and *ρ*_w,lit._ = 0.997g/ml is the literature value of the density of water at 23^*°*^C. The respective masses were measured using a high precision lab scale (Ohaus Pioneer PX124). The density of the liquid sample under study was then computed by

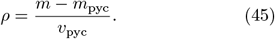

### C. Measurement uncertainties

The systematic uncertainties of the respective measurement devices under use were taken from the manuals and considered in all calculations together with the statistical uncertainties by employing Gaussian propagation of uncertainty as

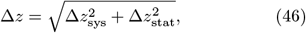

where *z* is an arbitrary observable.

### D. Data analysis

All data analysis, plotting and simulations were performed using custom scripts in `Wolfram Research, Inc., Mathematica, Version 12.2` [45].

## Supporting information

SI_MD_Estimation

## ACKNOWLEDGMENTS

We thank Abin Biswas, Giulia Zanini and Giuliano Scarcelli for the helpful and illuminating discussions on the topic.

