## Supplementary material for "Estimation of the mass density of biological matter from refractive index measurements": SI_MD_Estimation

### Supplemental Materials: Estimations of the mass density of biological matter from refractive index measurements

#### I. PARAMETER VALUES

In the following we provide the mean values ± standard deviations of the following quantities; relative volume fraction $x$, refractive index $n$, refraction per gram $R$ in $cm^3/g$, PSV $\theta$ in ml/g and RI increment $\alpha$ in ml/g of the different macro molecules employed throughout this study in Tab. SI. The RI and refraction per gram values of the lipids employed in the sections *Lipids and proteins in water* and *Larval zebrafish trunk tissue* were obtained from [46] (predicted data by `ACD/Labs Percepta Platform - PhysChem Module`, at $T = 20°C$ and a wavelength of $\lambda = 589\,nm$), from which we computed the PSV *via* the Lorentz-Lorenz relation (Eq. (10)). The associated standard deviation was then estimated *via* GPU. For all the computations presented in this study, we interpret a value that has a uncertainty attached to it as a normal distribution, where the standard deviation is given by said uncertainty. For values that were assumed to be precise, we assume a delta distribtuion.

#### II. PARTIAL AND APPARENT SPECIFIC VOLUMES AND THE REFRACTIVE INDEX INCREMENT

Based on derivations in [43], for a binary solution consistent of $N_1$ moles of solvent with molar volume $V_1$ and $N_2$ moles of solute, the total molar volume is given by

$$\begin{aligned} V &= N_1 V_1 + N_2 \tilde{\Theta}, \text{ or} \\ \tilde{\Theta} &= \frac{V - N_1 V_1}{N_2}, \end{aligned} \tag{S1}$$

where $\tilde{\Theta}$ denotes the apparent molar volume of the solute. This parameter accounts for the potential volume-in-additivity; in other words, it is a macroscopic quantity that describes the volume of the solute in dilution. The partial molar volume is given by

$$\Theta = \left(\frac{\partial V}{\partial N_2}\right)_{T,p,N_1} = N_2 \left(\frac{\partial \tilde{\Theta}}{\partial N_2}\right)_{T,p,N_1} + \tilde{\Theta}, \tag{S2}$$

and the apparent and partial specific volume can be expressed with the molar mass of the solute $M_2$ as $\tilde{\theta} = \tilde{\Theta}/M_2$ and $\theta = \Theta/M_2$, respectively.

##### A. Volume additivity

If the volume additivity is given, then

$$\begin{aligned} \Theta = V_2 = \tilde{\Theta} &= \frac{M_2}{\rho_2}, \text{ and} \\ \theta = \tilde{\theta} = 1/\rho_2 &= \frac{V_2 N_A}{M_2}. \end{aligned} \tag{S3}$$

##### B. Protein PSV and RI increment

As employed in [23, 34, 43, 50], the PSV and refraction per gram of a protein may be interpreted the weight average of all amino acids composing this protein as

$$\begin{aligned} \theta_p &= \frac{\sum_i V_{r,i} N_A}{\sum_i M_{r,i} + M_w}, \\ R_p &= \frac{\sum_i R^*_{r,i}}{\sum_i M_{r,i} + M_w}, \end{aligned} \tag{S4}$$

where $M_w = 18.01$ g/mol is the molecular mass of water [51, 52] and $N_A$ is the Avogadro constant.

TABLE SI: Rounded mean values ± standard deviations of relative volume fraction $x$, refractive index $n$, refraction per gram $R$ in $cm^3/g$, PSV $\theta$ in ml/g and RI increment $\alpha$ in ml/g of the different macro molecules employed throughout this study. Entries without footnote indicate assumptions or derived values (see main text for further information). Assumed to be precise values are stated without standard deviation.

| | $\overline{x} \pm \Delta x$ | $\overline{n} \pm \Delta n$ | $\overline{R} \pm \Delta R$ in $cm^3/g$ | $\overline{\theta} \pm \Delta\theta$ in ml/g | $\overline{\alpha} \pm \Delta\alpha$ in ml/g |
|---|---|---|---|---|---|
| Water | | 1.334 | | 1/0.997 | |
| Lipids and proteins in water | | | | | |
| Proteins (human proteome) | $(1-\overline{x}_{\text{lip}}) \pm \Delta x_{\text{lip}}$ | $1.602 \pm 0.007$[a] | $0.252 \pm 0.0032$[a] | $0.734 \pm 0.012$[a] | $0.196 \pm 0.004$[a] |
| Triolein (neutral lipid) | $\overline{x}_{\text{lip}} \pm \Delta x_{\text{lip}}$ | 1.477[b] | $0.30674 \pm 0.00034$[b] | $1.086 \pm 0.0012$ | $0.1552 \pm 0.0008$ |
| *Bovine* skim milk | | | | | |
| Proteins | 0.33[g] | | | | |
| Caseins | 0.20[c] | $1.596 \pm 0.007$[a] | $0.2531 \pm 0.0004$[a] | $0.744 \pm 0.007$[a] | $0.1957 \pm 0.0032$[a] |
| $\alpha_{S1}$-Casein | 0.42[c] | 1.600[a] | 0.253[a] | 0.740[a] | 0.198[a] |
| $\alpha_{S2}$-Casein | 0.11[c] | 1.602[a] | 0.253[a] | 0.737[a] | 0.199[a] |
| $\beta$-Casein | 0.35[c] | 1.588[a] | 0.254[a] | 0.753[a] | 0.192[a] |
| $\kappa$-Casein | 0.12[c] | 1.597[a] | 0.252[a] | 0.740[a] | 0.195[a] |
| Other proteins | 0.08[c] | $1.603 \pm 0.007$[a] | $0.2528 \pm 0.0030$[a] | $0.736 \pm 0.011$[a] | $0.198 \pm 0.004$[a] |
| Fat | 0.01[g] | 1.462[c] | 0.299 | 1.09[c] | 0.141 |
| Lactose | 0.52[g] | 1.582 | 0.187 | 0.562[c] | 0.140[c] |
| Ash | 0.06[g] | 1.647 | 0.196 | 0.541[c] | 0.170[c] |
| Intralipid solution | | | | | |
| Soybean oil | 0.89[g] | 1.474[h] | 0.31 | 1.09[h] | 0.15 |
| Glycerol | 0.07[g] | 1.474[h] | 0.22 | 0.79[h] | 0.11 |
| Lecithin | 0.05[g] | 1.459[i] | 0.27 | 0.97[j] | 0.12 |
| Larval zebrafish trunk tissue | | | | | |
| Proteins (trunk tissue) | $0.780 \pm 0.022$[d] | $1.602 \pm 0.011$[e] | $0.2519 \pm 0.0024$[e] | $0.734 \pm 0.009$[e] | $0.197 \pm 0.007$[e] |
| Lipids | $0.220 \pm 0.022$[d] | $1.473 \pm 0.016$ | $0.308 \pm 0.06$ | $1.100 \pm 0.023$ | $0.153 \pm 0.016$ |
| Triolein | 0.28[f] | 1.477[b] | $0.30674 \pm 0.00034$[b] | $1.0856 \pm 0.0012$ | $0.1552 \pm 0.0008$ |
| Palmitic acid | 0.24[f] | 1.454[b] | $0.3030 \pm 0.0012$[b] | $1.119 \pm 0.004$ | $0.1343 \pm 0.0029$ |
| Oleic acid | 0.20[f] | 1.467[b] | $0.3084 \pm 0.0011$[b] | $1.111 \pm 0.004$ | $0.1478 \pm 0.0026$ |
| Docosahexaenoic acid | 0.15[f] | 1.521[b] | $0.3224 \pm 0.0009$[b] | $1.0587 \pm 0.0030$ | $0.1980 \pm 0.0023$ |
| Stearic acid | 0.12[f] | 1.456[b] | $0.3058 \pm 0.0011$[b] | $1.125 \pm 0.004$ | $0.1373 \pm 0.0026$ |

[a] computed from AA sequences obtained from [30]
[b] obtained from [46]
[c] obtained from [27]
[g] obtained from the manufacturer
[h] obtained from [47]
[i] obtained from [48]
[j] obtained from [49]
[d] computed from values obtained from [40]
[e] computed from AA sequences obtained from [19] and [30]
[f] computed from values obtained from [41]

##### C. Errors with assuming volume additivity

- Errors in the PSV considerations
  - The following effects are stated in [43]
    - conformational protein formation from amino acids: $\sim 0.012$ ml/g
    - volume changes of proteins in solution:
      - swelling and dissolution: $\sim -5 \times 10^{-2}$ ml/g
      - thermal denaturation: $\sim \pm 10^{-2}$ ml/g
      - Coil-helix transition: $\sim +10^{-2}$ ml/g
      - Aggregation: $\sim +5 \times 10^{-3}$ ml/g

TABLE SII: Values of amino acid residues (AAR) employed for calculating the PSV $\theta$ and RI increment for Wiener approximation and volume additivity argument $\alpha_r^W$ and $\alpha_r^{VA}$, respectively. See text for further details. Values obtained from §[51, 52], †[34] (consensus average), ‡[53].

| AAR | $M_r^{\S}$ in g/mol | $V_r^{\dagger}$ in $10^{-3}$nm$^3$ | $\theta_r$ in ml/g | $R_r^{*\ddagger}$ in cm$^3$ | $\alpha_r^W$ in ml/g | $\alpha_r^{VA}$ in ml/g |
|---|---|---|---|---|---|---|
| Arg | 156.19 | $188 \pm 10$ | $0.73 \pm 0.04$ | $39.47 \pm 0.10$ | $0.194 \pm 0.016$ | $0.202 \pm 0.018$ |
| His | 137.14 | $156 \pm 6$ | $0.686 \pm 0.027$ | $34.62 \pm 0.15$ | $0.210 \pm 0.012$ | $0.221 \pm 0.014$ |
| Lys | 128.17 | $173 \pm 6$ | $0.811 \pm 0.028$ | $34.10 \pm 0.20$ | $0.184 \pm 0.012$ | $0.191 \pm 0.013$ |
| Asp | 115.09 | $115.4 \pm 2.2$ | $0.604 \pm 0.012$ | 26.06 | $0.193 \pm 0.005$ | $0.204 \pm 0.006$ |
| Glu | 129.12 | $141 \pm 5$ | $0.657 \pm 0.025$ | $30.07 \pm 0.15$ | $0.183 \pm 0.011$ | $0.192 \pm 0.012$ |
| Ser | 87.08 | $91.7 \pm 1.8$ | $0.634 \pm 0.012$ | $19.16 \pm 0.10$ | $0.168 \pm 0.006$ | $0.175 \pm 0.006$ |
| Thr | 101.10 | $118.3 \pm 2.3$ | $0.705 \pm 0.014$ | $23.82 \pm 0.10$ | $0.169 \pm 0.006$ | $0.175 \pm 0.007$ |
| Asn | 114.10 | $120 \pm 4$ | $0.634 \pm 0.022$ | $26.09 \pm 0.20$ | $0.185 \pm 0.010$ | $0.194 \pm 0.011$ |
| Gln | 128.13 | $145 \pm 5$ | $0.682 \pm 0.024$ | $30.37 \pm 0.20$ | $0.181 \pm 0.011$ | $0.189 \pm 0.012$ |
| 2Cys | 204.26 | $211 \pm 10$ | $0.621 \pm 0.029$ | 48.58 | $0.209 \pm 0.013$ | $0.221 \pm 0.016$ |
| Gly | 57.05 | $59.9 \pm 2.2$ | $0.632 \pm 0.023$ | $12.81 \pm 0.10$ | $0.177 \pm 0.011$ | $0.186 \pm 0.012$ |
| Pro | 97.12 | $123.2 \pm 1.8$ | $0.764 \pm 0.011$ | $23.74 \pm 0.10$ | $0.161 \pm 0.005$ | $0.167 \pm 0.005$ |
| Ala | 71.08 | $87.8 \pm 2.3$ | $0.744 \pm 0.019$ | $17.15 \pm 0.15$ | $0.164 \pm 0.009$ | $0.169 \pm 0.010$ |
| Ile | 113.16 | $166.1 \pm 3.4$ | $0.884 \pm 0.018$ | $31.87 \pm 0.20$ | $0.184 \pm 0.008$ | $0.191 \pm 0.009$ |
| Leu | 113.16 | $168 \pm 4$ | $0.894 \pm 0.023$ | $31.59 \pm 0.15$ | $0.176 \pm 0.010$ | $0.181 \pm 0.010$ |
| Met | 131.19 | $165.2 \pm 1.8$ | $0.758 \pm 0.008$ | $34.45 \pm 0.05$ | $0.199 \pm 0.004$ | $0.208 \pm 0.004$ |
| Phe | 147.18 | $190 \pm 7$ | $0.776 \pm 0.030$ | $42.21 \pm 0.15$ | $0.240 \pm 0.013$ | $0.253 \pm 0.016$ |
| Trp | 186.21 | $228 \pm 4$ | $0.737 \pm 0.012$ | $55.24 \pm 0.30$ | $0.278 \pm 0.007$ | $0.297 \pm 0.008$ |
| Tyr | 163.18 | $191 \pm 8$ | $0.706 \pm 0.030$ | 44.34 | $0.240 \pm 0.013$ | $0.255 \pm 0.016$ |
| Val | 99.13 | $139 \pm 4$ | $0.843 \pm 0.022$ | $26.73 \pm 0.13$ | $0.178 \pm 0.009$ | $0.184 \pm 0.010$ |

· Sol-gel transition: $\sim -10^{-4}$ ml/g

- Errors in the RI increment considerations
  - [50] found that the predicted RI increment values using the Wiener relation for different proteins are systematically higher than the experimental values. Further they found a linear relationship between predicted and experimental values. Further they propose a $\pi - \pi$ interaction term correction which slightly improves the agreement. We do not employ this correction here. However, we note that this discrepancy could also be partially resolved by not employing the Wiener equation but rather relying on the Biot equation, as well as a different choice of experimental values of $R_r^*$ for the amino acid residues.

##### D. Employed amino acid residues values

We omitted the contributions of selenocysteine since we could not obtain data on molar refractivity, mass density nor refractive index. For all other amino acid residues (AAR) the employed values are given in table SII.

#### III. EFFECT SIZES

##### A. Standardized effect size (SES)

One particular choice of effect size which stands in duality to the *Rayleigh* criterion, often referred to as standardized mean difference, or standardized effect size, is given by

$$d = \frac{|\mu_2 - \mu_1|}{\sqrt{(\sigma_1^2 + \sigma_2^2)/2}}, \tag{S5}$$

where $\mu_i$ and $\sigma_i$ are the mean and standard deviation of respective distributions. We note that for a skew-normal distribution $\mathcal{S}(\mu_0, \sigma_0, \eta_0)$, the mean and standard deviation is given by

$$\mu = \mu_0 + \sqrt{\frac{2}{\pi}} \frac{\sigma_0 \eta_0}{\sqrt{1+\eta_0^2}}, \; \sigma = \sigma_0 \sqrt{1 - \frac{2\eta_0^2}{\pi(1+\eta_0^2)}}, \tag{S6}$$

respectively.

##### B. Standardized quantile effect size (SQES)

Alternatively, when dealing with non-normal, or non-symmetric distributions, we could evaluate

$$d_{\mathrm{q}} = \frac{|q_2 - q_1|}{\sqrt{\left((q_{1+} - q_1)^2 + (q_2 - q_{2-})^2\right)/2}}, \tag{S7}$$

where $q_i$ are the 50-th quantiles, i.e., medians of the two distributions and $q_{i+}$ and $q_{i-}$ are the upper and lower bound of the $\gamma\%$ credible interval (CI) of the distribution $i$. For an arbitrary PDF of a parameter $\theta$ the $\gamma\%$ CI is implicitly defined by

$$\int_{q_{1-}}^{q_{1+}} \mathrm{d}\theta \mathcal{P}(\theta) = \gamma. \tag{S8}$$

A customary choice in biological sciences would be $\gamma = 68.3\%$.

In other words, $d_{\mathrm{q}}$ calculates the median difference, normalized by the 'pooled CI deviation'. In principle $d_{\mathrm{q}}$ is a generalization of the SES, however, its interpretation might not be as 'straightforward'.

##### C. Quantile absolute deviation (QAD)

Following the considerations in [42], another choice of effect size is given by the QAD

$$d_{\mathrm{QAD}} = \int_0^1 \mathrm{d}p \left|A^{-1}(p) - B^{-1}(p)\right|, \tag{S9}$$

where $A(t)$ and $B(t)$ are the respective CDFs of the two probability distributions $\mathcal{A}$ and $\mathcal{B}$ (see Fig. S1(c)) and $A^{-1}(p)$ and $B^{-1}(p)$ are their inverse functions, customarily called quantile function. A graphical representation of $\left|A^{-1}(p) - B^{-1}(p)\right|$ for two skew-normal distributions is given in Fig. S1(b).

##### D. Kolmogorov–Smirnov test statistic (KSTS)

Another choice of a bounded effect size is the KSTS, given by

$$d_{\mathrm{KS}} = \sup |A(t) - B(t)|, \tag{S10}$$

with $0 \leq d_{\mathrm{KS}} \leq 1$. See Fig. S1(c).

##### E. Quantile comparison effect size (QCES)

Introducing the QCES, denoted by $\Xi$, as proposed in [42]

$$\begin{aligned} D(A||B) &\equiv 2\int_0^1 \mathrm{d}p \left|A(B^{-1}(p)) - A(A^{-1}(p)\right| \\ &= 2\int_0^1 \mathrm{d}p \left|A(B^{-1}(p)) - p\right| \end{aligned} \tag{S11}$$

$$\Xi(A, B) \equiv \frac{1}{2}D(A||B) + \frac{1}{2}D(B||A). \tag{S12}$$

Here, $V_B^A(p) \equiv A(B^{-1}(p))$ is the so called *vertical quantile comparison function* (VQCF) and $V_B^A(p) - p$ provides a measure of divergence, which is not necessarily symmetric, i.e., $D(A||B) \neq D(B||A)$. However, the average of the two VQCFs, $\Xi$, is symmetric and therefore suited as an effect size, since the order of arguments should not matter. A graphical representation of $V_B^A(p)$ and $V_A^B(p)$ for two skew-normal distributions is given in Fig. S1(d).

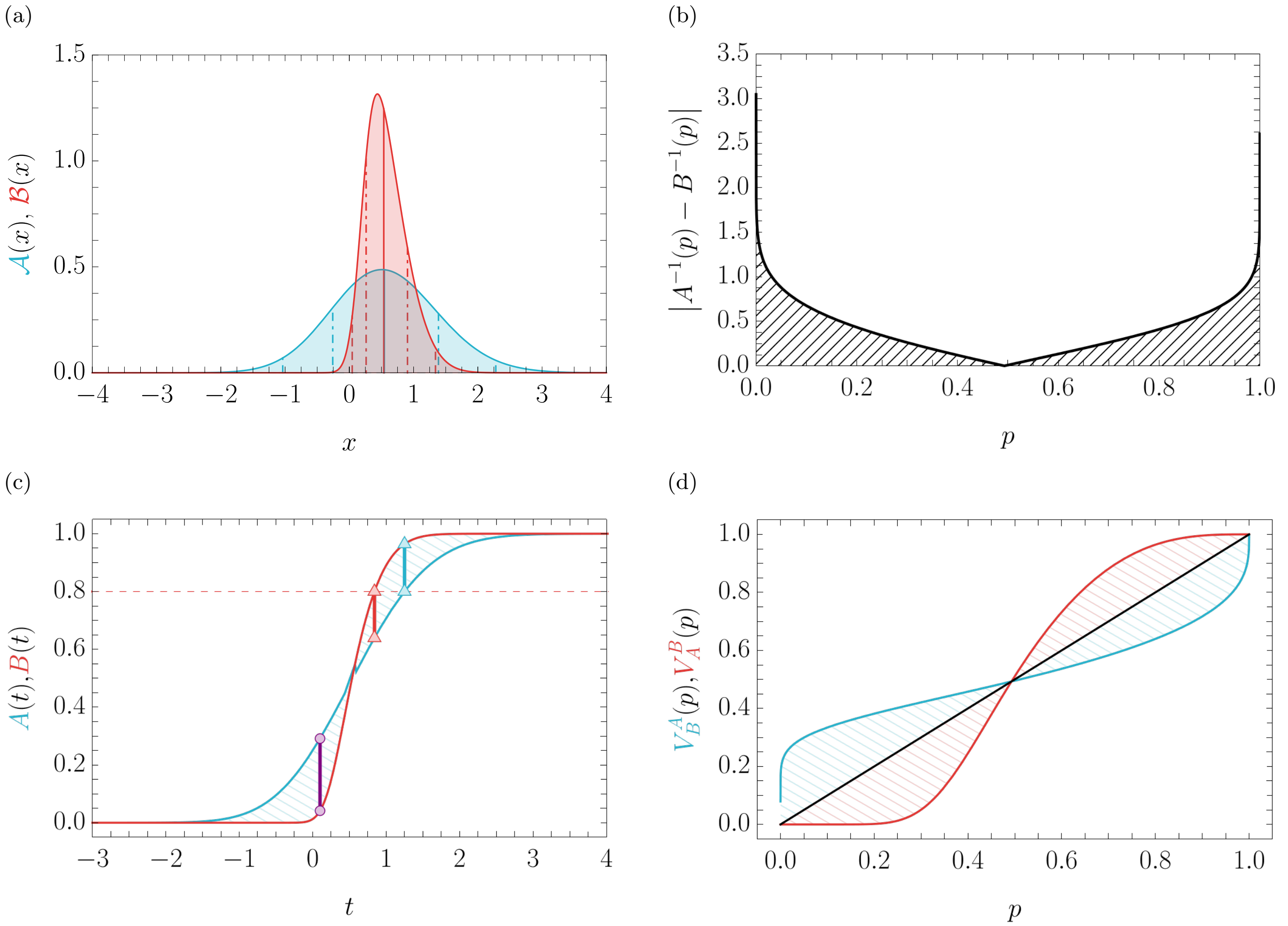


FIG. S1: (a): PDFs $\mathcal{A}(x) = \mathcal{S}(0, 1, 1; x)$ (blue) and $\mathcal{B}(x) = \mathcal{S}(0.2, 0.5, 3; x)$ (red), (b): $\left|A^{-1}(p) - B^{-1}(p)\right|$ and $d_{\text{QAD}}$ indicated by the hatched filling, (c): CDFs $A(t)$ (blue) and $B(t)$ (red) and respective derived quantities: $d_{\text{KS}}$ (purple) and $(A(B^{-1}(0.8)), 0.8)$ (blue) and $(B(A^{-1}(0.8)), 0.8)$ (red), (d): VQCFs $V_B^A(p)$ (blue) and $V_A^B(p)$ (red). The hatched filling indicates $D(A||B)$ and $D(B||A)$, respectively.

#### IV. MEAN AND STANDARD DEVIATION OF NORMAL MIXTURE DISTRIBUTIONS

Let's consider a one dimensional mixture distribution of two Normal distributions,

$$\mathcal{P}(\mu_1, \mu_2, \sigma_1, \sigma_2; x) = w\mathcal{N}(\mu_1, \sigma_1; x) + (1 - w)\mathcal{N}(\mu_2, \sigma_2; x), \tag{S13}$$

with $0 \leq w \leq 1$. The mean and standard deviation of this mixture distribtion is given by

$$\begin{aligned} \mu' &= w\mu_1 + (1 - w)\mu_2, \\ \sigma' &= \sqrt{\sigma_2^2 + w\left(\mu_1^2 - 2\mu_1\mu_2 + \mu_2^2 + \sigma_1^2 - \sigma_2^2\right) - w^2\left(\mu_1^2 - \mu_2^2\right)^2}, \end{aligned} \tag{S14}$$

respectively. For the special case of $\sigma_1 = \sigma_2 = \sigma$ we obtain for the standard deviation of the mixture distribution

$$\sigma' = \sqrt{\sigma^2 - \left((w - 1)\, w\, (\mu_1 - \mu_2)^2\right)} > \sigma, \tag{S15}$$

for all $\mu_1 \neq \mu_2$ and $0 \leq w \leq 1$, which is maximized for $w = 0.5$.

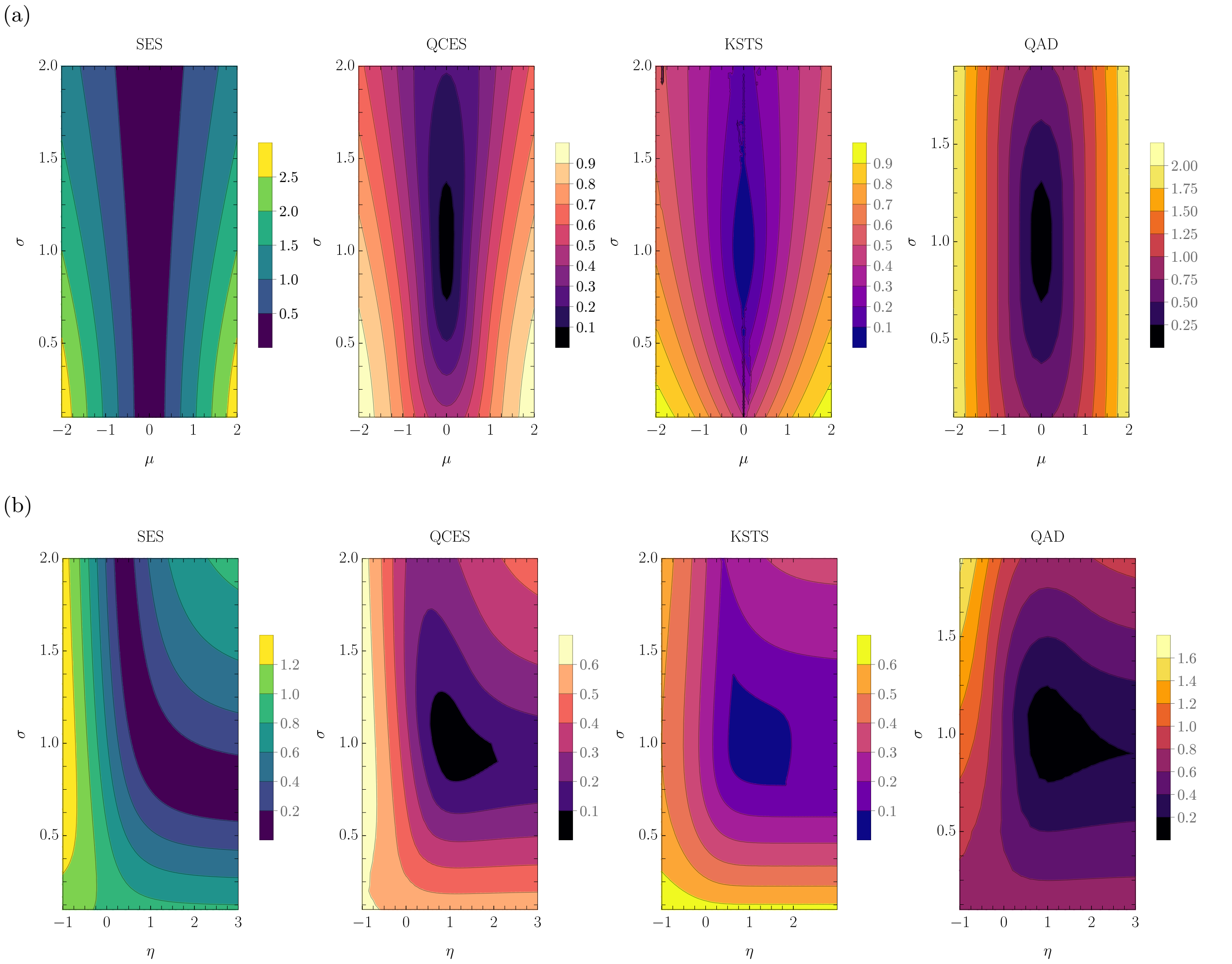


FIG. S2: Different significance estimates SES (Eq. (S5)), QCES (Eq. (S11)), KSTS (Eq. (S10)) and QAD (Eq. (S9)) for the explicit PDFs (a): $\mathcal{A}(x) = \mathcal{N}(0,1;x)$, $\mathcal{B}(x) = \mathcal{N}(\mu,\sigma;x)$ and (b): $\mathcal{A}(x) = \mathcal{S}(0,1,1;x)$ and $\mathcal{B}(x) = \mathcal{S}(0,\sigma,\eta;x)$

#### V. EFFECTIVE PARAMETERS

From the mixing rule of the solute density

$$\rho_{\mathrm{s}} = \sum_{i=1}^{N_{\mathrm{s}}} x_{i+1}\rho_{i+1}, \tag{S16}$$

we obtain that the effective PSV is

$$\theta_{\mathrm{eff}} = 1/\rho_{\mathrm{s}} = \left(\sum_{i=1}^{N_{\mathrm{s}}} \frac{x_{i+1}}{\theta_{i+1}}\right)^{-1} = \sum_{i=1}^{N_{\mathrm{s}}} y_{i+1}\theta_{i+1} = \sum_j \frac{N_j m_j}{m_{\mathrm{s}}}\theta_j, \tag{S17}$$

where $N_\mathrm{s}$ is the nuber of solute voxelinos per voxel and $j$ denotes the different types of solute molecules. The effective RI increment for the Biot equation is then

$$\begin{aligned}\alpha_\mathrm{eff} &= \frac{n-n_1}{c} = \frac{n_1\left(1-c\sum_{i=1}^{N_\mathrm{s}} y_{i+1}\theta_{i+1}\right)+c\sum_{i=1}^{N_\mathrm{s}} y_{i+1}\theta_{i+1}n_{i+1}-n_1}{c}\\ &= n_1\sum_{i=1}^{N_\mathrm{s}} y_{i+1}\theta_{i+1}-\sum_{i=1}^{N_\mathrm{s}} n_{i+1}y_{i+1}\theta_{i+1}\\ &= \sum_{i=1}^{N_\mathrm{s}}(n_1-n_{i+1})y_{i+1}\theta_{i+1}\\ &= \sum_{i=1}^{N_\mathrm{s}} y_{i+1}\alpha_{i+1} = \sum_j \frac{N_j m_j}{m_\mathrm{s}}\alpha_j = \theta_\mathrm{eff}\left(n_\mathrm{s}-n_1\right).\end{aligned} \tag{S18}$$

#### VI. UNCERTAINTIES OF THE EFFECTIVE PARAMETERS FOR LIPIDS AND PROTEINS IN WATER

The uncertainties of the effective RI increment and the PSV, introduced in Eq. (30) can be computed as follows. The terms $\Delta\alpha^0_\mathrm{eff}$ and $\Delta\theta^0_\mathrm{eff}$ refer to the standard deviations of the mixture distribution, as given in Eq. (S14).

The uncertainties associated of the effective PSV to deviations in the relative lipid volume fraction $\Delta x_\mathrm{lip}$, can be evaluated by employing Eq. (24), from which we obtain that

$$\frac{\partial\overline{\theta}_\mathrm{eff}}{\partial\overline{x}_\mathrm{lip}} = \frac{\overline{\theta}_\mathrm{p}\overline{\theta}_\mathrm{lip}\left(\overline{\theta}_\mathrm{lip}-\overline{\theta}_\mathrm{p}\right)}{\left(\left(1-\overline{x}_\mathrm{lip}\right)\overline{\theta}_\mathrm{lip}+\overline{x}_\mathrm{lip}\overline{\theta}_\mathrm{p}\right)^2}. \tag{S19}$$

We may write the effective RI increment, using Eq. (24), as

$$\overline{\alpha}_\mathrm{eff} = \overline{\theta}_\mathrm{eff}\left(\overline{n}_\mathrm{s}-n_1\right) = \overline{\theta}_\mathrm{eff}\left(\left(1-\overline{x}_\mathrm{lip}\right)\overline{n}_\mathrm{p}+\overline{x}_\mathrm{lip}\overline{n}_\mathrm{lip}-n_1\right). \tag{S20}$$

Hence, we have

$$\frac{\partial\overline{\alpha}_\mathrm{eff}}{\partial\overline{x}_\mathrm{lip}} = \overline{\theta}_\mathrm{eff}\left(\overline{n}_\mathrm{lip}-\overline{n}_\mathrm{p}\right)+\frac{\partial\overline{\theta}_\mathrm{eff}}{\partial\overline{x}_\mathrm{lip}}\frac{\overline{\alpha}_\mathrm{eff}}{\overline{\theta}_\mathrm{eff}}. \tag{S21}$$

#### VII. INTERPRETATION OF A VOXELINO VOLUME

In crystallography it is customary to define the the molar volume of a unit cell by

$$V_\mathrm{m} = \frac{v_\mathrm{cell}}{Z}, \tag{S22}$$

where $Z$ is the number of formula units per crystallographic unit cell. In our case, the volume of the unit cell is a volume of a voxelino $v_0$. Consequently, number of formula units may be connected to the PSV of the voxelino by

$$Z = \frac{N_\mathrm{A}v_0}{\theta M}, \tag{S23}$$

where $M$ is the molar mass of the consituent present in the voxelino.

#### VIII. REFRACTIVE INDEX VALUES OF LARVAL ZEBRAFISH TRUNK TISSUE AT 96 HPF FROM [19]

The RI values of larval zebrafish trunk tissue at 96 hpf employed in this study are not explicitly stated in [19], hence we list them here.

TABLE SIII: Refractive indices $n$ and according standard deviations $\Delta n$ of the trunk tissue of $N = 20$ zebrafish larvae at 96 hpf from [19].

| $N$ | 1 | 2 | 3 | 4 | 5 | 6 | 7 | 8 | 9 | 10 |
|---|---|---|---|---|---|---|---|---|---|---|
| $n$ | 1.36501 | 1.36524 | 1.36691 | 1.36679 | 1.36556 | 1.36251 | 1.36319 | 1.36098 | 1.36112 | 1.36342 |
| $\Delta n$ | 0.00225 | 0.00369 | 0.00307 | 0.00315 | 0.00268 | 0.00151 | 0.00122 | 0.00210 | 0.00241 | 0.00229 |
| $N$ | 11 | 12 | 13 | 14 | 15 | 16 | 17 | 18 | 19 | 20 |
| $n$ | 1.36763 | 1.36604 | 1.36673 | 1.36670 | 1.36550 | 1.36640 | 1.36633 | 1.36632 | 1.36662 | 1.36693 |
| $\Delta n$ | 0.00182 | 0.00205 | 0.00233 | 0.00211 | 0.00219 | 0.00238 | 0.00220 | 0.00172 | 0.00130 | 0.00238 |

#### IX. ESTIMATIONS OF $x_{\mathrm{lip}}$ AND $\varphi_1$ OF LARVAL ZEBRAFISH AT 96 HPF

Based on the measurements of [40], where they determined the wet mass $m_{\mathrm{tot}}$, dry mass $m_{\mathrm{dry}}$, protein mass $m_{\mathrm{p}}$ and the lipid mass $m_{\mathrm{lip}}$ of larval zebrafish at 96 hpf, we define the relative lipid mass fraction $y_{\mathrm{lip}} \equiv m_{\mathrm{lip}}/(m_{\mathrm{lip}} + m_{\mathrm{p}})$ and consider the ratio

$$\frac{y_{\mathrm{lip}}}{y_{\mathrm{p}}} = \frac{m_{\mathrm{lip}}}{m_{\mathrm{p}}} = \frac{\theta_{\mathrm{p}}^{\mathrm{eff}} x_{\mathrm{lip}}}{\theta_{\mathrm{lip}}^{\mathrm{eff}}(1 - x_{\mathrm{lip}})}, \tag{S24}$$

from which we obtain

$$x_{\mathrm{lip}} = \frac{m_{\mathrm{lip}}\theta_{\mathrm{lip}}^{\mathrm{eff}}}{m_{\mathrm{lip}}\theta_{\mathrm{lip}}^{\mathrm{eff}} + m_{\mathrm{p}}\theta_{\mathrm{p}}^{\mathrm{eff}}}, \tag{S25}$$

and analogously

$$\varphi_1 = \frac{(m_{\mathrm{tot}} - m_{\mathrm{dry}})}{m_{\mathrm{tot}} + m_{\mathrm{dry}}\left(\rho_1\theta_{\mathrm{dry}}^{\mathrm{eff}} - 1\right)}. \tag{S26}$$

Based on this dry mass composition we estimate the distributions of $\theta_{\mathrm{p}}^{\mathrm{eff}}$ and $\theta_{\mathrm{lip}}^{\mathrm{eff}}$ by running the MC simulation of the extended mixture model with $x_{\mathrm{lip}} = 0$ and $x_{\mathrm{lip}} = 1$, respectively. We may now obtain the distribution of $x_{\mathrm{lip}}$ by a MC sampling approach, considering the distributions of all individual parameters (see Fig. S3(a)). Next, we compute $\theta_{\mathrm{dry}}^{\mathrm{eff}}$ by repeating the procedure described above for $x_{\mathrm{lip}}$ following the distribution determined previously. With that we obtain the distribution of $\varphi_1$, as shown in Fig. S3(b).

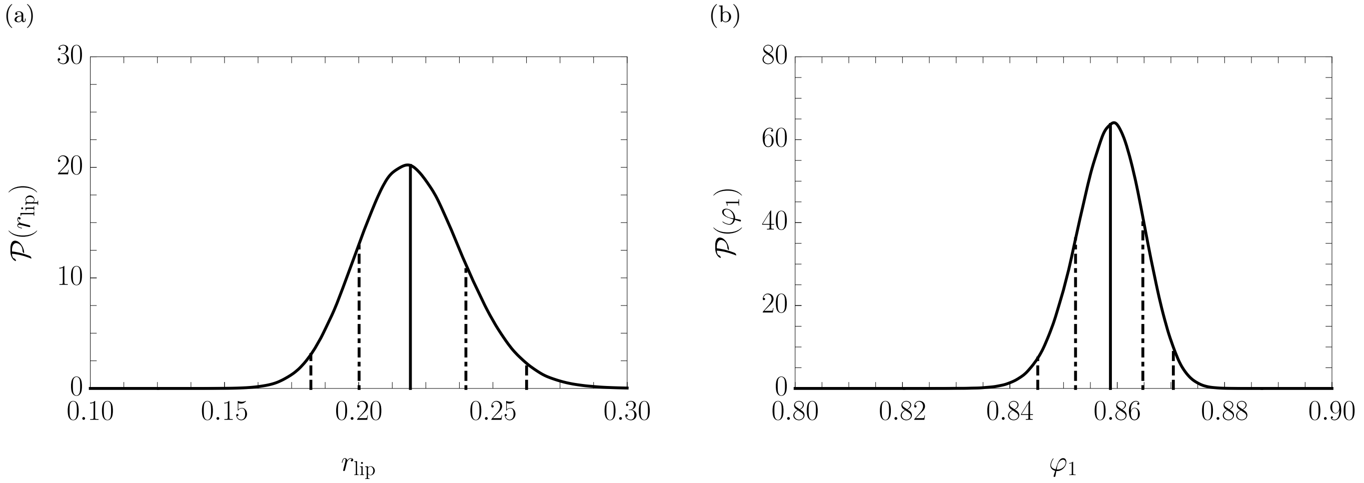


FIG. S3: Estimated PDFs of the relative lipid volume fraction $x_{\mathrm{lip}}$ ((a)) and the water volume fraction $\varphi_1$ ((b)) for the spinal cord tissue of the zebrafish larva 96 hpf.

#### X. RELATIVE UNCERTAINTY OF THE RI FOR DIFFERENT MATERIAL PROPERTIES

The dependence of the relative RI deviation on the deviation of the relative lipid volume fraction $\Delta x_{\mathrm{lip}}$ for different deviations of the water volume fraction $\Delta\varphi_1$, as pointed out in the main text, can be qualitatively understood with Fig. S4.

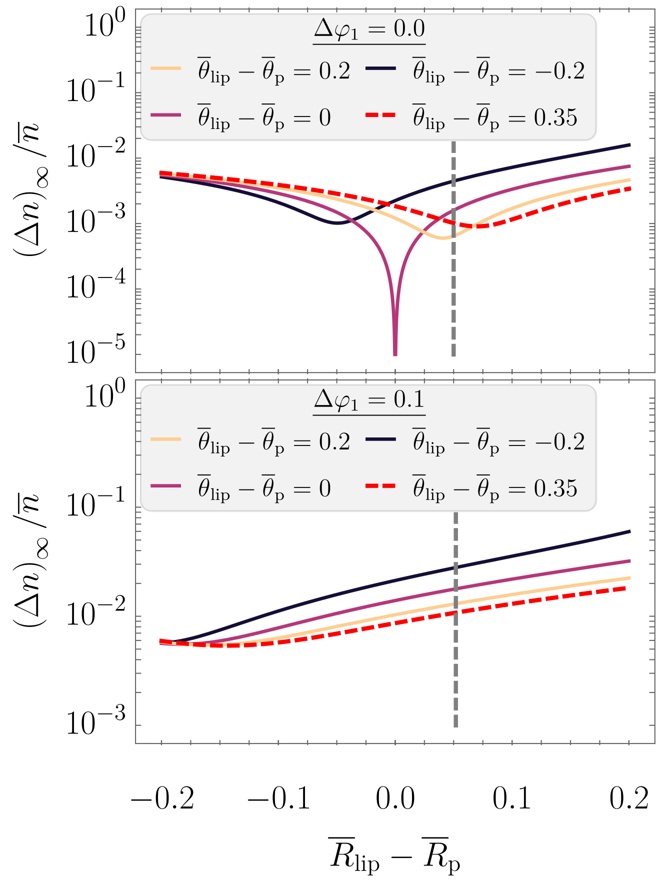


FIG. S4: Asymptotic relative deviation of the RI $(\Delta n)_\infty/\overline{n}$ for different potential differences of lipid and protein PSVs $\overline{\theta}_{\mathrm{lip}} - \overline{\theta}_{\mathrm{p}}$ in dependence of the difference of lipid and protein refractions per gram $\overline{R}_{\mathrm{lip}} - \overline{R}_{\mathrm{p}}$ for the cases of a non-fluctuating water volume fraction $\Delta\varphi_1 = 0$ (top) and a fluctuating water volume fraction $\Delta\varphi_1 = 0.1$ (bottom).

#### XI. STRATEGIES FOR ESTIMATING THE MD FOR CERTAIN EXPERIMENTAL PARADIGMS

In Fig. S5, we outline possible strategies for estimating the MD, given certain experimental insights. In the following, we use the same abbreviations as in the main text, namely,

- RI = refractive index,
- (S)RS = (stimulated) Raman spectroscopy,
- MS = mass spectrometetry.

We want to point out that the estimation process is heavily dependent on identifying relevant solute constituents of the sample, as well as their PSVs, RIs and/or refractions per gram. For the latter, secondary data bases, such as ChemSpider [46], are invaluable. Furthermore, although RI measurements are not necessary to predict $\rho(\delta n)$, as outlined in the main text, comparing predicted and measured RI distributions gives necessary insight on the validity of the prediction.

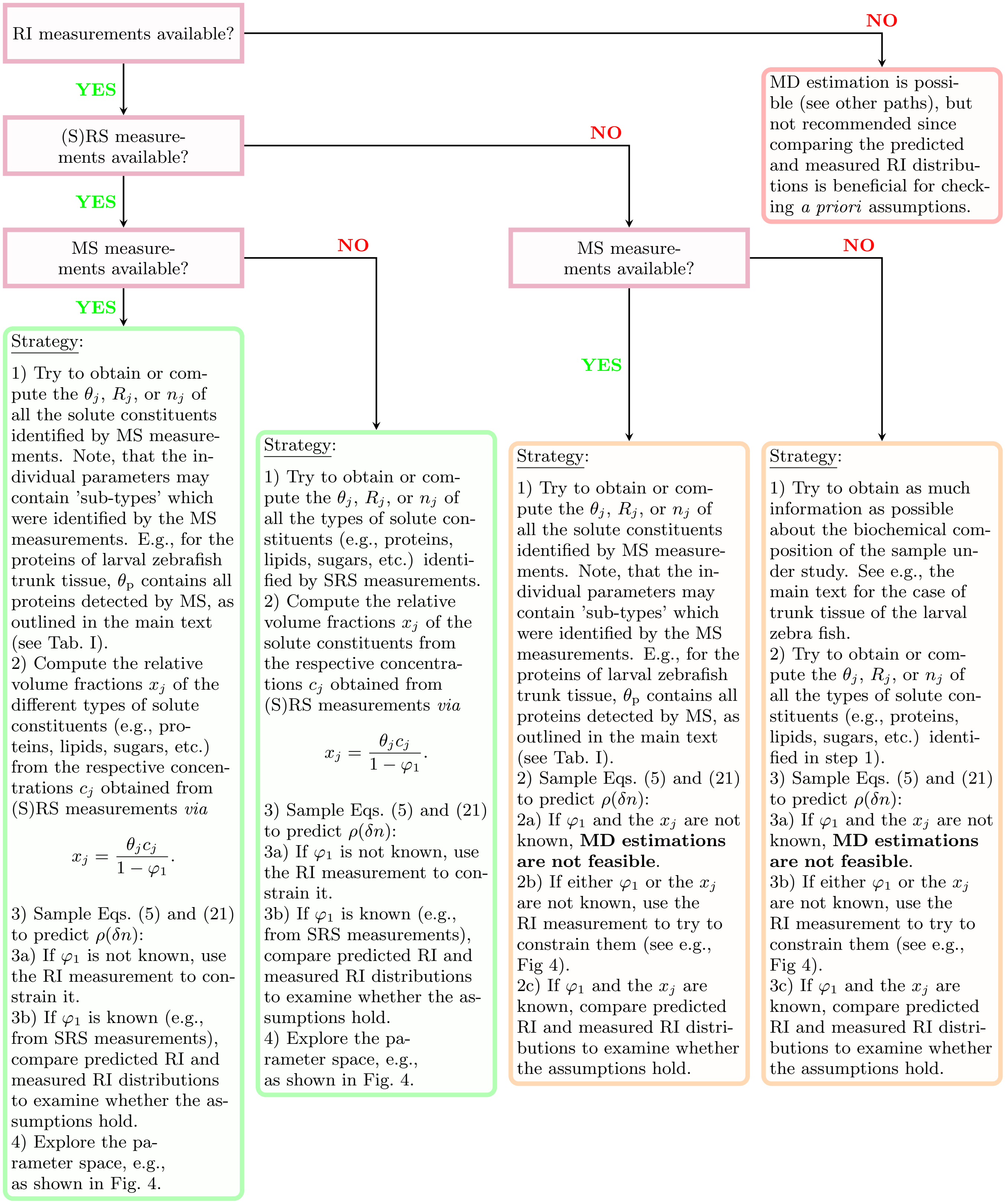


FIG. S5: Flowchart of mass density estimation approaches outlined in this study, based on different experimental paradigms. The abbreviations and symbols are as defined in the main text.
